# Large-scale map of RNA binding protein interactomes across the mRNA life-cycle

**DOI:** 10.1101/2023.06.08.544225

**Authors:** Lena Street, Katherine Rothamel, Kristopher Brannan, Wenhao Jin, Benjamin Bokor, Kevin Dong, Kevin Rhine, Assael Madrigal, Norah Al-Azzam, Jenny Kim Kim, Yanzhe Ma, Ahmed Abdou, Erica Wolin, Ella Doron-Mandel, Joshua Ahdout, Mayuresh Mujumdar, Marko Jovanovic, Gene W Yeo

## Abstract

Messenger RNAs (mRNAs) interact with RNA-binding proteins (RBPs) in diverse ribonucleoprotein complexes (RNPs) during distinct life-cycle stages for their processing and maturation. While substantial attention has focused on understanding RNA regulation by assigning proteins, particularly RBPs, to specific RNA substrates, there has been considerably less exploration leveraging protein-protein interaction (PPI) methodologies to identify and study the role of proteins in mRNA life-cycle stages. To address this gap, we generated an RNA-aware RBP-centric PPI map across the mRNA life-cycle by immunopurification (IP-MS) of ∼100 endogenous RBPs across the life-cycle in the presence or absence of RNase, augmented by size exclusion chromatography (SEC-MS). Aside from confirming 8,700 known and discovering 20,359 novel interactions between 1125 proteins, we determined that 73% of our IP interactions are regulated by the presence of RNA. Our PPI data enables us to link proteins to life-cycle stage functions, highlighting that nearly half of the proteins participate in at least two distinct stages. We show that one of the most highly interconnected proteins, ERH, engages in multiple RNA processes, including via interactions with nuclear speckles and the mRNA export machinery. We also demonstrate that the spliceosomal protein SNRNP200 participates in distinct stress granule-associated RNPs and occupies different RNA target regions in the cytoplasm during stress. Our comprehensive RBP-focused PPI network is a novel resource for identifying multi-stage RBPs and exploring RBP complexes in RNA maturation.

**HIGHLIGHTS:**

- An RBP-centric RNA-aware PPI network focuses on the mRNA life-cycle in human cells
- Prey-prey correlation analysis assigns prey proteins to life-cycle stages, of which 536 proteins (half of the network) interact with multiple steps
- ERH is highly connected to multiple RNPs to affect nuclear speckle organization and mRNA export
- Splicing factor SNRNP200 interacts with stress granule proteins and has distinct RNA occupancy in the cytoplasm

## INTRODUCTION

RNA molecules associate with thousands of RNA-binding proteins (RBPs) to construct functional ribonucleoprotein complexes (RNPs) (Baltz et al. 2012; Dreyfuss, Kim, and Kataoka 2002; Beckmann et al. 2015). These RNPs are dynamic across RNA life-cycle steps, with RBPs rearranging and exchanging both RNA substrates and protein interactors to orchestrate enzymatic processing, subcellular localization, mRNA translation, and degradation of mRNAs. Mass spectrometry approaches and computational prediction models (Brannan et al. 2016; Gerstberger et al. 2014; Hentze et al. 2018; Castello et al. 2016; Queiroz et al. 2019; Zhao et al. 2014; Bressin et al. 2019; X. Zhang and Liu 2017; Beckmann, Castello, and Medenbach 2016; Ben-Bassat, Chor, and Orenstein 2018; Ghanbari and Ohler 2020) estimate >5000 RNA-interacting proteins in the human genome, with the majority of these having unknown RNA regulatory and biological functions.

RNPs are believed to be functionally modular, with distinct RBP combinations driving RNP localization across subcellular compartments and forming dynamic, RNP-concentrated granular structures (Achsel and Bagni 2016; Lunde, Moore, and Varani 2007). Nuclear speckles and stress granules are two such highly dynamic structures that serve as hubs for numerous protein-RNA interactions (Guillén-Boixet et al. 2020; Sanders et al. 2020; Faber, Nadav-Eliyahu, and Shav-Tal 2022; Spector and Lamond 2011; Strom and Brangwynne 2019; Protter and Parker 2016). Nuclear speckles, rich in splicing factors and other RNA processing proteins, facilitate pre-mRNA splicing and other RNA maturation events. Stress granules, in contrast, form in the cytoplasm to modulate mRNA translation in response to various stresses. Aberrant formation or persistence of such RNP granules has been implicated in the pathogenesis of many human diseases, including neurodegeneration and cancer (Guillén-Boixet et al. 2020; Shorter 2019; McSwiggen et al. 2019; Marmor-Kollet et al. 2020; Seiler et al. 2018; Desterro, Bak-Gordon, and Carmo-Fonseca 2020) Defining the RNP composition at the interchange of such granules across life-cycle steps could elucidate disease mechanisms.

Current methods primarily focused on identifying the RNA targets of individual RBPs using crosslinking and immunoprecipitation (CLIP) followed by subsequent sequencing of the targets (C. Zhang and Darnell 2011; Hafner et al. 2010; Zarnegar et al. 2016; König et al. 2010). ENCODE (Van Nostrand, Pratt, et al. 2020; Blue et al. 2022; Van Nostrand et al. 2016; Van Nostrand, Freese, et al. 2020) has performed enhanced CLIP (eCLIP) on ∼300 RBPs and has strictly validated nearly a thousand endogenous IP-grade antibodies (Sundararaman et al. 2016). These CLIP methods are powerful for obtaining individual RBP binding profiles in isolation, but they are blind to RNP-complex membership and dynamics. Because RBPs can be found in multiple RNP complexes, contain diverse RNA-binding domain (RBD) combinations, and shuttle between subcellular locales, they likely have multiple functions within and across mRNA life-cycle stages that can be uncovered by RBP-focused protein-protein interaction (PPI) networks (García-Mauriño et al. 2017). RBP PPIs modulate RNA-binding affinity, specificity, and functionality, therefore to characterize multifunctional RBPs, it is critical to define both RBP-targets and RBP protein-interactors (Lunde, Moore, and Varani 2007; Bleichert and Baserga 2010).

Systematic large-scale efforts to characterize PPIs across cell types, such as OpenCell (Cho et al. 2022), Bio-ID (Roux et al. 2018), BioPlex (Huttlin et al. 2021, 2017, 2015), and the Human Interactome (Hein et al. 2015) rely on fusion of affinity tags and bait over-expression, both of which can produce non-physiological interactomes and confound analysis of the stoichiometries between interaction partners. Moreover, while some of these networks do have considerable RBP bait representation, they fail to assess the RNA dependency of these interactions, and therefore cannot distinguish if interactions between two proteins are direct or RNA-mediated. RBP-focused attempts to delineate RBP PPI networks also have similar limitations, including lack of RNA-aware interaction information and use of exogenous tags to immunoprecipitate or localize the RBPs (Garriga-Canut et al. 2020; Kwon et al. 2013). For example, the yeast two-hybrid screen (rec-Y2H) (Lang et al. 2021) generated RNA-independent pairwise binding maps of 978 human RBPs, however the incorporation of nuclear localization signals in each protein likely resulted in some spurious interactions.

As an orthogonal method to study PPIs, size exclusion chromatography followed by mass spectrometry (SEC-MS), can identify interactions by measuring the co-elution of protein pairs or complexes across size exclusion fractions (Kristensen, Gsponer, and Foster 2012; Kirkwood et al. 2013; Skinnider et al. 2021; Heusel et al. 2019; Rosenberger et al. 2020; Fossati et al. 2021; Bludau et al. 2020). SEC-MS complements IP-MS methods as it is not biased by bait selection and can therefore build an interaction network that detects prey-prey interactions. Moreover, SEC-MS eliminates reliance on antibodies that may exhibit preferences for specific bait conformations or exhibit off-target binding. The utility of co-elution approaches to studying RBPs has been previously demonstrated. R-DeeP applied sucrose density gradients in the presence and absence of RNase to define the RNA-dependent proteome (Caudron-Herger et al. 2019), and was primarily used to identify proteins that interact with RNA, through a shift in the elution profile between conditions. Similarly, DIF-FRAC (Mallam et al. 2019) systemically characterized known protein complexes as RNA-associated in HEK293T and mouse embryonic stem cells by applying SEC-MS in RNase treatment versus control conditions. While SEC is able to complement IP-MS interactomes and has been shown to be useful in studying RBPs, SEC is limited in its ability to capture substoichiometric interactions or proteins that are lowly expressed. Previous work has shown that substoichiometric interactions are key links within a larger network, connecting stable modules that show interaction stoichiometries around 1:1 ratios (Hein et al. 2015). These factors point towards the importance of capturing the full range of a protein’s interactions, including those that are less abundant or less stable, when trying to understand regulation of processes on a cellular level. Thus, an ideal approach would contain experimental information from both IP-MS and SEC-MS.

We used such a combined proteomics approach to gain insight into the protein-RNA networks of RBPs across the full mRNA life-cycle. We developed an RNase-coupled proteomics strategy to identify protein partners of known RBPs (e.g., our baits) in HEK293XT cells. Examining PPIs in the presence and absence of RNase allowed us to capture both RNA-mediated and direct PPIs. This strategy discovers new RBPs by exploiting that RBPs often form complexes with one another, and also identifies RBP complexes and individual RBPs that may concurrently bind to the same RNA substrates. Thus, these data gave us greater insight into how individual RBPs can form multiple functional complexes to mediate gene expression. Our PPI network was centered around 92 RBPs found across several cellular compartments and from every major stage of mRNA maturation (transcription, splicing, translation, etc.). From this analysis, we generated a network of 1125 proteins, forming 29,545 interactions, which we classified into specific interaction categories and compared to other established PPI networks. To uncover novel RNP complexes, RBP function, and mRNA life-cycle nexuses, we conducted multiple prey-centric and betweenness centrality analyses. These analyses revealed numerous multifunctional RBPs, leading to the detailed characterization of the two highly connected proteins ERH and SNRNP200. Our network analyses, in combination with eCLIP and functional assays, describe novel cellular compartment-specific functions and binding profiles of these two proteins.

## RESULTS

### An RBP-centric, RNA-aware PPI network captures novel RBP interactions and functions across the mRNA life-cycle

Using the mRNA life-cycle as a guide to map PPIs of RBPs that are either RNA-dependent or - independent (e.g., direct), we selected ∼100 RBPs with major roles in RNA processing at one or more mRNA life-cycle steps, namely transcription, splicing, modification, 3’ end processing, nuclear export, localization, translation, and degradation (Figure 1A, Supplemental Table 1). The availability of ENCODE-validated, immunopurification (IP)-grade antibodies that effectively recognize these endogenous RBPs was an important selection criterion for these bait RBPs (Van Nostrand, Freese, et al. 2020; Sundararaman et al. 2016). Upon initial creation of the PPI network based on the first round of baits (Figure 1A), we discovered prey proteins ERH, DLD, DLST and RUVBL2 that were central to the network or involved in unexpected interactions and also utilized these proteins as baits in the full network.

**Figure 1.**
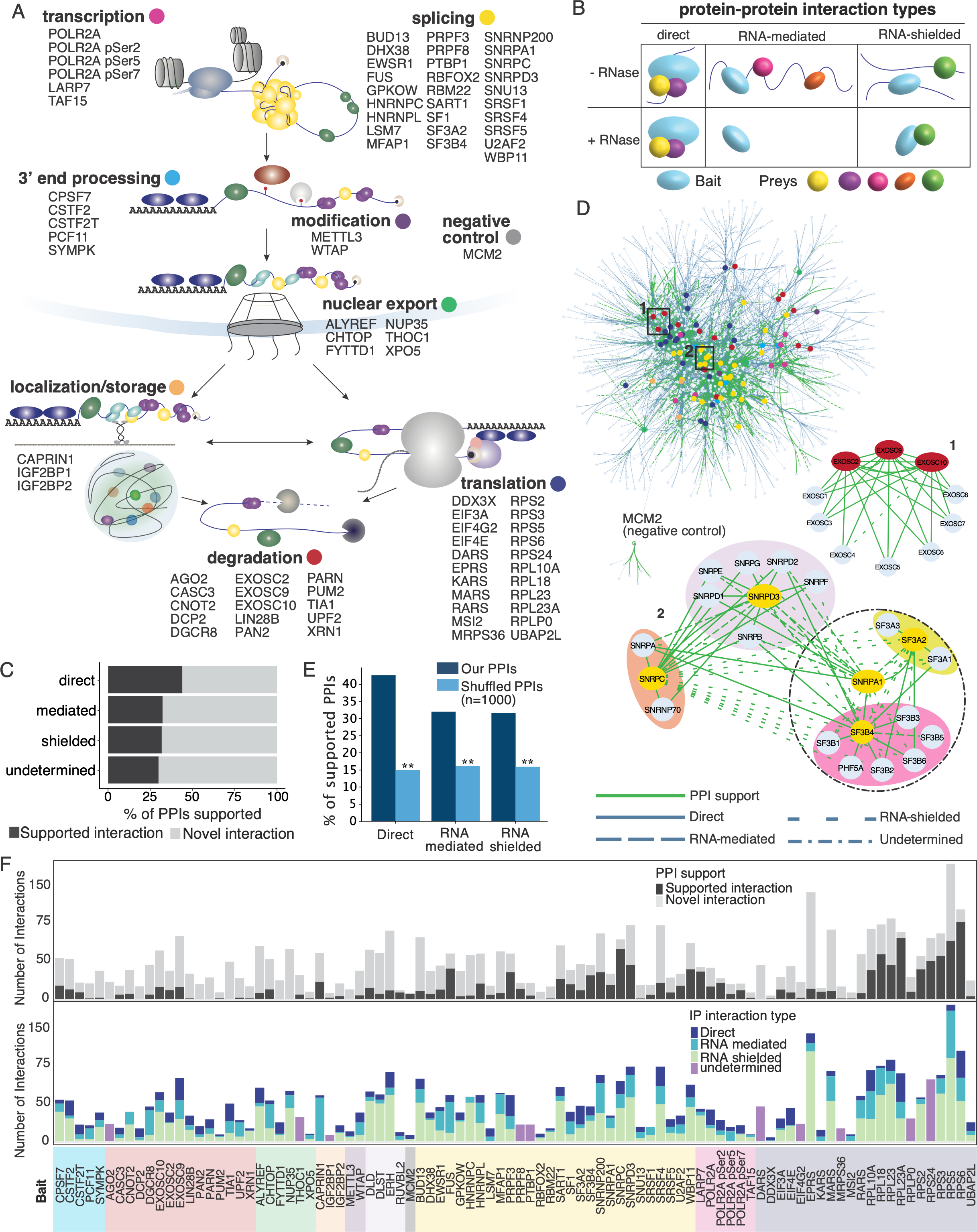
Endogenous immunopurification followed by LC-MS/MS captures and quantifies the RNA-dependent or independent interactome of RNA-binding-proteins across the mRNA lifecycle. A. mRNA life-cycle assignments of baits targeted by IP-MS to build the RBP centric protein-protein interactome. Almost all bait RBPs have evidence for functions within more than one life-cycle stage so primary assignment was based on literature review. B. Types of RNA-independent and dependent interactions that can be found in the network as a result of RNase treatment. Direct interactions are found with and without RNase treatment. RNA-mediated are found without RNase and lost upon RNase treatment. RNA-shielded are found with RNase treatment but not without. One possible mechanism behind RNA shielded interactions is depicted, whereby proteins may be found in more than one binding state depending on the presence of RNA. C. The IP-MS network as viewed in cytoscape with bundled edge layout. Bait nodes are colored by their life-cycle stage assignments in A. The mRNA exosome (1) and U1, U2, and Sm core spliceosome complex (2, Sm core [purple], U1 specific proteins [orange], U2 specific proteins: SF3a complex [yellow], SF3b complex [pink]) interactions are zoomed in to show how interaction types behave within the network. D. Percent of novel and supported interactions in the network for each interaction type. E. Percent of supported interactions in the network compared to randomly shuffled interactions of all the proteins in our network for n=1000 tests, p-value < 0.001. F. The number, types, and percent support of interactions for each bait sorted by mRNA life-cycle stage assignment.

To identify both the direct and RNA-mediated protein interactors (PIs) for each bait, IPs were performed in triplicate in the presence and absence of RNase. Samples were multiplexed by TMT11 (Thermo Scientific; for higher throughput and quantitative accuracy), measured by liquid chromatography-coupled tandem mass spectrometry (LC-MS/MS), and raw data was searched using SpectroMine (Biognosys) (Supplemental Figure 1A). After batch correction and QC (see methods), 92 bait proteins remained in the dataset. We classified interactions (Figure 1B) as follows: (1) direct PIs were successfully immunopurified by a bait regardless of the presence or absence of RNA (i.e. detected in both with and without RNase conditions). (2) RNA-mediated PIs were immunopurified only in the presence of RNA (without RNAse condition); these RNA-mediated interactions do not necessarily have to result from RBPs binding RNA but include PIs that depend on the presence of RNA. (3) RNA-shielded PIs were immunopurified only in the absence of RNA (with RNase condition); and (4) undetermined PIs were those where the IP for a specific bait failed across one of the RNase conditions and therefore we could not determine the RNA dependence of the interactions (Supplemental Table 1). We expected that the majority of interactors would fall into the first two categories of direct and RNA-mediated interactions. However, almost half our interactions fell into the third category, RNA-shielded interactions, in which the interactions were only detected in the absence of RNA. We found that 88.7% (1820/2052) of the preys found in shielded interactions were most closely associated with the splicing and translation hubs of the network. Furthermore 21% of the shielded interactions were found in CORUM in large, well-defined complexes such as the ribosome and spliceosome (Supplemental Figure 1B). We also found that RNA-shielded PIs are supported in the literature at levels similar to RNA-mediated and undetermined interactions (Figure 1C). These results indicate that large RNP complexes may be difficult to IP in strict RNase-minus conditions, however upon partial RNA digestion, the bait proteins may be in smaller sub-complexes or have their antibody-binding residues exposed such that they can be enriched by IP. Finally, 10 antibodies only enriched their baits in either with-RNase or without-RNase conditions; therefore, these bait-specific interactions were classified as undetermined.

Our resulting IP-MS-generated RBP PPI network contains 1125 distinct proteins (nodes) with 4351 interactions (edges) (Figure 1D, Supplemental Table 2). We discovered 886 Direct, 1108 RNA-mediated, 2052 RNA-shielded, and 305 undetermined interactions, of which 497 (56.1%), 751 (67.8%), 1401 (68.3%) and 214 (70.2%) are novel, respectively, when compared to PPI databases and large-scale interactome studies (Giurgiu et al. 2019; Tsitsiridis et al. 2023; Cho et al. 2022; Kuksa et al. 2020; Calderone, Castagnoli, and Cesareni 2013; Lang et al. 2021; Roux et al. 2018; Rolland et al. 2014) (Figure 1C). We observed that 34.2% (1488/4351) of all the interactions were previously identified (Supplemental Figure 1B), which is within the range of other recent large-scale interactomes (BioPlex 3.0 - 9.3% (Huttlin et al. 2021), BioPlex 2.0 - 13% (Huttlin et al. 2017), Human Interactome 16% (Hein et al. 2015), 62% OpenCell (Cho et al. 2022), 73% (Havugimana et al. 2022), and our interactome adds RNA-dependence information to these interactions. To ensure that the support for our PPIs is not artificially inflated due to high connectivity, we compared our interactions to randomly shuffled interactions within the network. Our direct interactions showed nearly three times more support compared to random interactions (resampling test, n = 1000, p-value < 1e^-3^ for all interaction types, Figure 1E), and RNA-mediated and shielded interactions demonstrated approximately two times more. We further validated our findings by performing the same test using only CORUM PPIs (Tsitsiridis et al. 2023; Giurgiu et al. 2019), a benchmark dataset, and found that direct interactions had over four times more support than random pairs, while RNA-mediated and shielded interactions had just under three times more (p-values < 0.001, Supplemental Figure 1C). Finally, as we focused the interactome around the mRNA life-cycle, we expected that many of the prey proteins in the network would be predicted to be RBPs. Using the RBP predictor HydRa (Jin et al. 2023) we see that the prey proteins in the network have significantly higher RNA-binding prediction scores compared to our measured HEK293XT total proteome, with over ∼75% of preys predicted to be RBPs (Supplemental Figure 1D, K-S test p-value < 2.2e^-16^). These results highlight the high quality of our network, which not only aligns with the literature and other large-scale networks, but also uncovers a significant number of new interactions throughout the mRNA life-cycle.

Regarding connectivity, 59.7% of preys interact with more than one bait and 48.5% interact with baits of more than one life-cycle stage (Supplemental Figure 1E). At the center of the network are two hubs of splicing and translation baits with the RBP baits across the remainder of the life-cycle distributed more uniformly (Figure 1D). When we compared the life-cycle and interaction type distribution of the total interactome to the previously reported interactions (Supplemental Figure 1F) we found that direct interactions had higher levels of literature support than other types of interactions across the life-cycle. Moreover, our RNase treatment approach helped us gain previously uncharacterized but important information about the role of RNA in influencing PPIs throughout the mRNA life-cycle. As a negative control for the connectivity of our network, we included minichromosome maintenance complex component II (MCM2), a protein involved in genome replication. Since we expected the mRNA life-cycle interactome to be highly interconnected, we wanted to ensure that RBP interactors were not identified simply because of high local concentrations, especially in cellular compartments such as the nucleus. Indeed, the MCM2 interactome hub is distinct from the rest of the network, unlike the life-cycle baits, which form highly interconnected hubs (Figure 1D). MCM2 forms direct interactions with its canonical interactors in the MCM complex (MCM3-7), a histone chaperone, DNAJC9, which has been seen to interact with MCM2 (Hammond et al. 2021), and SUPT16H, a nucleosome remodeling factor; MCM2’s one connection to the rest of the network is via an RNA-shielded interaction with the nuclear receptor protein NCOA5.

We next asked if our interactome and the RNA-dependency information confirmed our expectations for well-studied complexes (Figure 1D). As expected for stable complexes that form without an RNA scaffold, the members of the mRNA exosome were connected by direct interactions. In contrast, the U1 and U2 spliceosome complexes are connected by a mixture of direct, RNA-mediated, and RNA-shielded interactions. Within the larger complex, the SF3a, SF3b, and Sm-snRNA core complexes as well as the U1 specific proteins had direct intra-complex connections. The inter-complex connections were a mix of direct, mediated, and shielded interactions as would be expected for a larger complex that assembles in the presence of RNA. Overall, our network recapitulates known PPIs and also captures undiscovered RBP interactions, providing a comprehensive map of RBP PPIs and their RNA-dependency across the mRNA life-cycle (Figure 1F).

### SEC-MS elucidates complex membership within the IP-MS interactome

IP-MS experiments capture PPIs with high sensitivity and at a range of stoichiometric ratios; however, they are limited to capturing only bait-prey interactions, and are unable to group complex membership amongst sets of prey proteins. Specifically, if a bait protein is a member of two different complexes, IP-MS can capture the members of those complexes but cannot disentangle which preys are members of one complex versus the other because there is no information about prey-prey interactions. To address this limitation, we performed size-exclusion chromatography followed by mass spectrometry (SEC-MS) in HEK293XT cells (Figure 2A, Supplemental Figure 2A). Cells were lysed under IP conditions and fractionated on a Yarra-4000 column (Phenomenex). SEC fractions were measured individually by LC-MS/MS and the resulting data was analyzed using CCprofiler (Heusel et al. 2019; Bludau et al. 2020). We focused our analysis on discovering the composition of protein complexes in our IP-MS interactome, by testing for statistically significant co-elution of interacting proteins as compared to a decoy interaction set. The strength of this approach is that it fills in our targeted IP-MS network by incorporating both IP-MS interactions with orthogonal SEC-MS support and prey-prey interactions, providing a more complete understanding of the complex interactions involved in the mRNA life-cycle.

**Figure 2.**
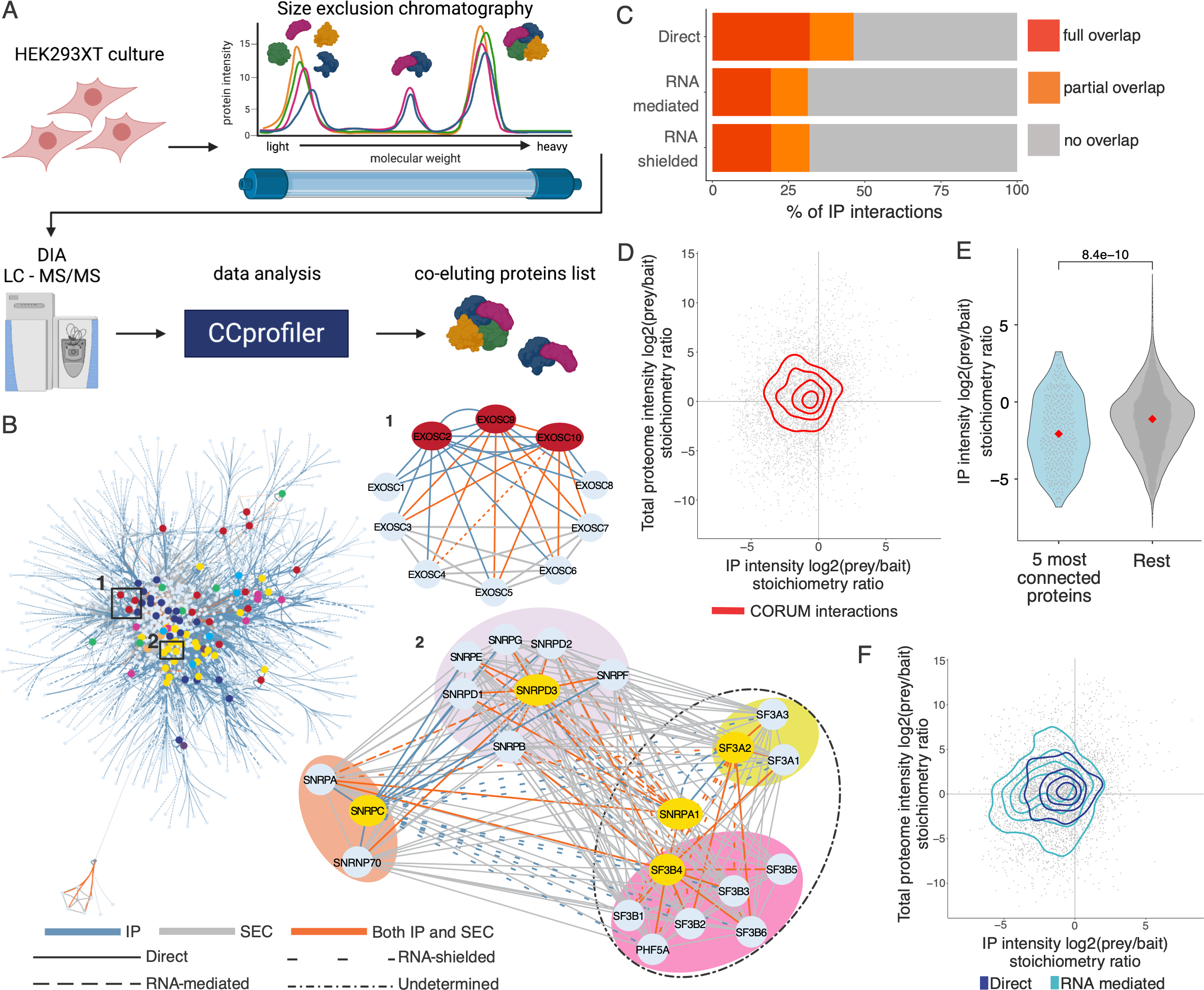
Size Exclusion Chromatography followed by LC-MS/MS identifies prey-prey interactions and provides orthogonal support for the IP-MS network, creating a comprehensive network with modular structure. A. Schematic of the SEC-MS experimental workflow. Size exclusion chromatography was performed in the absence of RNase followed by liquid chromatography and mass spectrometry. The data was searched to identify co-eluting proteins within the IP-MS PPI network to determine SEC-MS support of IP-MS interactions and assign prey-prey interactions. B. The combined IP-MS and SEC-MS interactome. Bait nodes are colored by life-cycle stage assignments as in Figure 1A and the mRNA exosome and Sm core, U1 and U2 spliceosome complexes are zoomed-in as in Figure 1C, showing how the SEC-MS data augments the IP-MS data. C. Co-localization of interacting proteins in the network based on the proteins’ Human Protein Atlas localization assignments. Interacting proteins where 100% of their assignments were shared are labeled as “full overlap”, where only some were shared are “partial overlap”, and where there were no shared localization assignments are “no overlap”. D. Stoichiometric ratios of interactions found in the highly curated CORUM database. The log2 ratios of the intensities of the prey over bait in the IP-MS data is plotted against the log2 intensity ratios of the prey over bait in total proteome data. E. The stoichiometric ratios in the IP-MS data for the 5 most connected proteins in the network are significantly lower (adjusted p-value Student’s t-test) than the ratios of the rest of the network. Medians are plotted in red. F. Stoichiometric ratios of direct and RNA-mediated interactions are plotted as in D.

The resulting SEC-MS interactome identified 26,946 interactions between the 1125 proteins in the IP-MS network, of which 1752 interactions (40.3% of the IP-MS interactome) provide orthogonal confirmation for IP-MS interactions and 25,194 are prey-prey interactions. Of the SEC-MS interactions, 8005 (29.7%) have been previously described in the literature and 18,941 are novel (Supplemental Figure 2B). We performed the same resampling test used on the IP-MS network to test if the SEC interactions have more literature support than random pairs of proteins in the network and found that the SEC interactions are about 2 times more supported than expected by chance at a p-value < 1e^-3^ (n = 1000) and about 3 times more supported in CORUM than expected by chance (p-value < 1e^-3^, n = 1000) (Supplemental Figure 2C and D). Based on these results we were confident that the SEC-MS data would enhance the IP-MS interactome so we combined the IP-MS and SEC-MS networks into a final interactome of 29,545 interactions (of which 8746, 29.6% confirm previous findings) between 1125 proteins.

SEC was particularly helpful in strengthening parts of the interactome with a lower IP-MS bait density. For example, the U1 and U2 spliceosomal interactome becomes much more highly connected upon addition of the SEC data (Figure 2B), enabling greater resolution of how the sub-complexes involved in nuclear mRNA processing interact. Conversely, three subunits of the mRNA exosome were selected as IP baits and thus the complex already has good coverage in the IP-MS network; thus, while SEC was able to strengthen the resolution of the complex by identifying prey-prey interactions between the subunits and provide orthogonal support for many of the IP-MS interactions, the complex was already quite well-defined in the IP-MS network (Figure 2B).

We next looked more closely at the overlap between the IP-MS and SEC-MS interactomes. Of the 1752 interactions found in both IP and SEC, 750 (42.8%) were literature-supported compared to 28.4% of IP-only interactions and 29.2% of SEC-only interactions (Supplemental Figure 2B), indicating that as expected, interactions found by orthogonal methods are more likely to have been previously found by other approaches. We next asked if we could use the SEC data to improve our understanding of the RNA-dependence of our IP-MS interactions. We hypothesized that a large percentage of the RNA-shielded interactions found by IP-MS were characterized as shielded due to limitations of the IP method and not because they only exist in the absence of RNA; if our interpretation is correct, those interactions should be captured in the RNase-minus SEC network at the same rate as direct or RNA-mediated interactions since SEC does not have the same size or epitope availability limitations as IP.

Instead of seeing similar percentages of PIs supported in SEC across interaction types, we actually found that the percent of RNA-shielded interactions was greater than any other type; 46.2% of interactions in the IP-MS network were shielded and that number increased to 53% of interactions found in both IP and SEC (compared to 21.7% vs 16% for direct and 25% vs 23.8% for RNA-mediated for IP vs IP+SEC, respectively) (Supplemental Figure 2E). In total, 29.9% of IP-defined direct interactions, 37.8% of RNA-mediated, and 45.3% of RNA-shielded interactions were recapitulated in SEC. These results strengthen the hypothesis that a large percentage of the RNA-shielded interactions are actually found in the presence of RNA but limitations of the IP method alone prevent their accurate identification. Thus, combining both strategies enhances our understanding of the complexes involved in the mRNA life-cycle. The general division of the network into nuclear and cytoplasmic compartments that was observed in the IP-MS network is preserved as are the general positions of the baits within the network. The interactome is supported at levels comparable to other recent large-scale interactomes, while additionally identifying novel interactions, providing RNA-dependency information for the IP-MS interactions, identifying prey-prey interactions that improve complex resolution, and providing orthogonal confirmation of interactions found in both IP-MS and SEC-MS.

### Interacting proteins show significant subcellular co-localization and life-cycle stages are connected via substoichiometric interactions

We next evaluated the RBP PPI network for network-level properties to reveal more about how RBPs function within the mRNA life-cycle. An important feature of RBPs is that even those proteins with assigned canonical functions or mRNA life-cycle step associations often play roles in other functions and parts of the cell. As protein localization is a reasonable proxy for function and is directly related to proteins’ ability to interact, we wondered to what extent our interactome recapitulated previous subcellular localizations. We found that our interactions had significantly greater co-localization assignments in the Human Protein Atlas (Thul et al. 2017) compared to random interactions between proteins in the interactome and were co-localized at similar levels to other large-scale datasets, such as BioPlex 3.0 (Huttlin et al. 2021) (Supplemental Figure 2F). Analysis of how previously-reported co-localizations were distributed across different types of interactions revealed that direct interactions had the greatest overlap with 46.4% of direct interactions having full or partial overlap of assigned bait and prey localizations, compared to 31.4% of RNA-mediated, and 32% of RNA-shielded interactions (Figure 2C). These levels are comparable to the 34% of BioPlex 3.0 interactions with co-localized annotations.

We next asked if the general network structure could give us insights into the flow of molecules throughout the life-cycle. When we observe how our bait proteins with assigned life-cycle stages distribute within the full-scale network at low resolution, we see a high density of interactions in the center of the network, which is primarily composed of distinct hubs of splicing and translation proteins with the other stages more generally distributed throughout the network (Figure 2B, Supplemental Table 3). At deeper resolution we observe smaller, highly intra-connected hubs that recapitulate well-described complexes such as the transcription-export (TREX) or the cleavage stimulation factor (CSTF) complexes (Supplemental Figure 2G). However, there are a large number of proteins that cannot primarily be assigned to one hub but rather form connections between hubs. To be able to gain more general insight into how interactome proteins behave when they are in hubs compared to forming connections between hubs, we turned to the stoichiometric information present in our IP-MS data.

Stoichiometric information from IP-MS experiments can be used to understand network structure (Cho et al. 2022; Hein et al. 2015). Specifically, the relative intensities of the baits and preys in the MS data reflect proxies of their interaction stoichiometries. Here, we estimated the total proteome abundance of the proteins in the interactome by LC-MS/MS measurements of total cell lysates so that we could ensure that proteins were not showing low stoichiometric ratios simply due to lower total abundance of one of the interactors. When we plotted the stoichiometric ratios of the interactions from the IP-MS data against the ratios from the total protein abundances we found that, as expected, interactions in CORUM, which we use as a proxy for well-defined stable complexes, were centered around 1:1 stoichiometries (Figure 2D). We next looked at our most connected proteins (see below and Figure 3B) and their interactors to ask if we also see that proteins responsible for connecting distinct network hubs form substoichiometric interactions. We found that the top 5 most connected proteins in our network have significantly lower interaction stoichiometries than the rest of the network (Figure 2E). Additionally, when we looked at the subsequent most connected proteins (top 10 and 15) we found that they also had significantly lower, but increasing median, interaction stoichiometries (Supplemental Figure 2H). These results indicate that highly connected proteins at the center of our interactome may make more frequent but less stable interactions to help facilitate movement of mRNA and proteins throughout the mRNA life-cycle.

**Figure 3.**
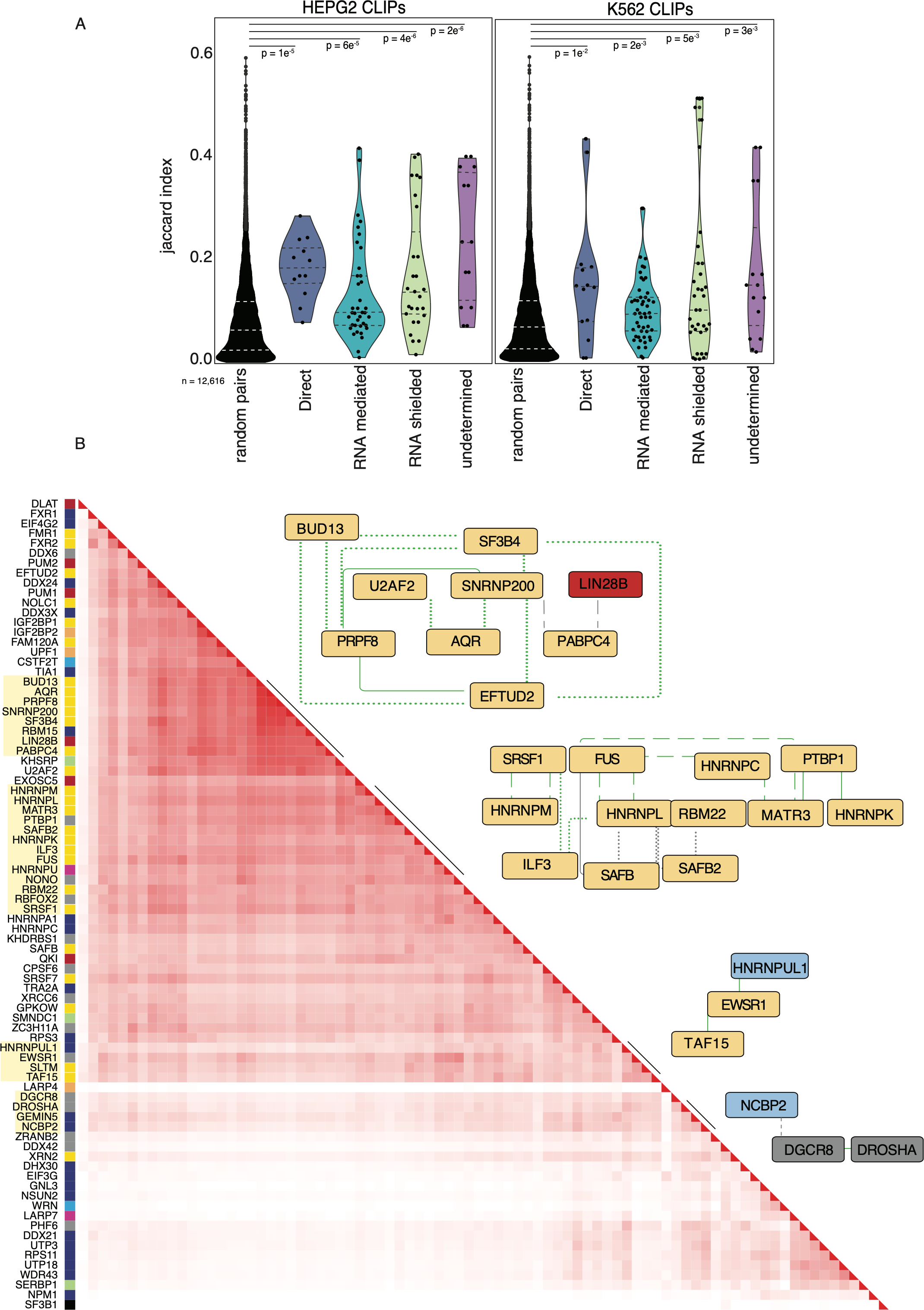
ENCORE data support RNA-aware PPI network. A. Violin plot of the Jaccard index scores across interaction types and cell types from ENCODE data. Each dot represents a pair of RBP interactions. Dashed lines represent the median and the first and third quartiles. B. Hierarchical clustering heatmap of the jaccard indices for protein pairs in K562 cells. Recapitulated network interactions are highlighted next to the corresponding cytoscape plot.

We next queried whether the RNA-dependency of interactions correlates with differences in interaction stoichiometries. Direct interactions overlap well with CORUM interactions, which is expected as we see the greatest overlap of CORUM interactions with direct interactions (Figure 2F, Supplemental Figure 2I). RNA-mediated interactions have lower and more varied stoichiometric ratios compared to direct interactions, and a wider total proteome abundance ratio range - as would be expected of proteins whose expression and degradation are not co-regulated as seen for many proteins that are found in stable complexes (Eisenberg et al. 2018; Taggart et al. 2020; McShane et al. 2016) (Figure 2F, Supplemental F2I). RNA-shielded interactions have interaction stoichiometries between those of direct and RNA-mediated, consistent with the large number of shielded interactions that map to the ribosome and spliceosome (Supplemental Figure 2I), meaning that the shielded distribution tended towards 1:1 ratios due to many of those interactions. The differences in ratios between interaction types indicate that direct interactions are likely more stable and represent distinct enzymatic hubs, while RNA-mediated interactions may be more transient and represent mRNPs functional in the progression of mRNA processing. Taken together, these results show that although RBPs can form discrete subnetworks, the RBP interactome behaves similarly to the cellular proteome interactome. Notably, RNP hubs with well-defined functions outline the mRNA life-cycle stages and are connected by proteins with substoichiometric interactions.

### Known complexes scaffold the RBP network and respond predictably to RNase treatment

As indicated by our analysis of the network structure, known complexes form hubs within the network and provide a scaffold upon which connecting protein interactions form. 62.7% (5487/8746) of the known interactions in our interactome are found in CORUM complexes. Importantly we see that complexes with known higher-order structures, such as RNA polymerase II and the mRNA exosome (Supplemental Figure 2G), consist primarily of direct interactions (4.5:1 and 26:0 direct:mediated, respectively) and thus these complexes can persist whether or not RNA is present. Conversely, complexes known to be scaffolded by RNA, such as the spliceosomal subcomplexes and the ribosome, are connected by a larger proportion of RNA-mediated interactions (1.1:1 and 1.2:1 direct:mediated, respectively; Supplemental Figure 2G). It is useful to note that while distinct cellular compartments such as nucleus and cytoplasm are often thought of as independent cellular regions, the reality is likely that they are more connected. When we visualize only those interactions from the network that are described in CORUM, we see a large number of interactions connecting the spliceosomal and ribosomal hubs (Supplemental Figure 2G). These interactions could have several explanations including that they may result from ribosomal protein shuttling into the nucleolus, persistence of RBPs primarily annotated as splicing factors on mRNA that is exported out of the nucleus, or from experimental artifacts.

### Transcriptome-wide binding data validates network structure

To test the hypothesis that interacting RBPs are more likely to bind to similar transcripts, we performed a co-occurrence analysis using the Jaccard index, leveraging available ENCODE eCLIP data (Van Nostrand, Pratt, et al. 2020; Van Nostrand et al. 2016) (Supplemental Table 4). The Jaccard index measures the intersection of two sets of data divided by the sample size of each of the datasets. 72 and 63 RBPs within our network have eCLIP data in HepG2 and K562 cell-lines, respectively. We reasoned that RBPs that form complexes and interact in any manner would have a higher probability of binding the same transcripts as RBPs that do not interact. In both cell types, all interaction types in our network have significantly higher Jaccard indexes than random pairs of RBPs in the ENCODE data (Wilcox; p-values (direct PPIs) = 1e^-5^, 1e^-2^; p-values (RNA-mediated PPIs) = 6e^-5^, 2e^-3^; p-values (RNA-shielded PPIs) = 4e^-6^, 5e^-3^; p-values (undetermined PPIs) = 2e^-6^, 3e^-3^ in HepG2 and K562 cells, respectively) (Figure 3A). We observe that for HepG2 and K562 cells undetermined (0.222 and 0.14) and direct (0.172 and 0.14) interactions have the highest median Jaccard index, respectively. In contrast, RNA-shielded (0.09 and 0.13) and RNA-mediated (0.09 and 0.09) interactions have lower Jaccard indices than direct and undetermined but still significantly higher indices than random pairs of RBPs. Together, these data indicate that, although all PPI interaction types are co-binding their RNA targets more significantly than expected by chance, direct and shielded PPIs are more likely to bind similar binding sites on transcripts than proteins that interact in an RNA-mediated manner.

We then performed hierarchical clustering on the Jaccard indices to test if we could recapitulate PPIs within our network (Figure 3B). The analysis revealed several large clusters that mimicked distinct interactions from our network. In our network, the splicing factor HNRNPL interacts directly with HNRNPC and MATR3 and interacts in an RNA-dependent manner with ILF3 and CSTF2. MATR3, HNRNPC, and HNRNPL have been previously described as forming a multimeric splicing complex (Damianov et al. 2016). The core spliceosome components ETUD2, PRPF8, BUD13, SNRNP200, and SF3B4 also clustered, highlighting the precision of our network and that interactions can be recapitulated using binding site data. This also supports our hypothesis that RBPs that interact bind RNAs as a complex.

### Prey-centric analysis unveils life-cycle and protein complex organization

Our list of preys includes a substantial proportion of known and predicted RBPs (Supplemental Figure 1F). However, many of these candidate RBPs have yet to be functionally linked to specific RNA life-cycle stages. To investigate the functional relationships within the network and generate hypotheses about the functions of these uncharacterized preys, we examined the distribution of life-cycle steps associated with each prey (Figure 4A). This analysis involved assigning each prey to the major life-cycle role of the baits with which it interacts (as depicted in Figure 1), except for a few baits (ERH, DLD, DLST and RUVBL2) that have not been characterized in the mRNA life-cycle and were labeled as unknown. By linking the preys to specific RNA life-cycle steps based on their interactions with the baits, we were able to identify clusters of functionally similar preys and assign them to specific life-cycle steps (Supplemental Figure 3A). Interestingly, a significant portion (50%) of preys were assigned to multiple life-cycle steps (Supplemental Table 5). Additionally, a subset of preys that are associated with splicing and translation also interact with preys from other mRNA life-cycle steps, suggesting that RBPs associated with splicing and translation in particular might have roles outside of their known functions. This analysis provides valuable interaction signatures that serve as starting points for determining the functional roles of previously uncharacterized prey proteins in our network.

**Figure 4.**
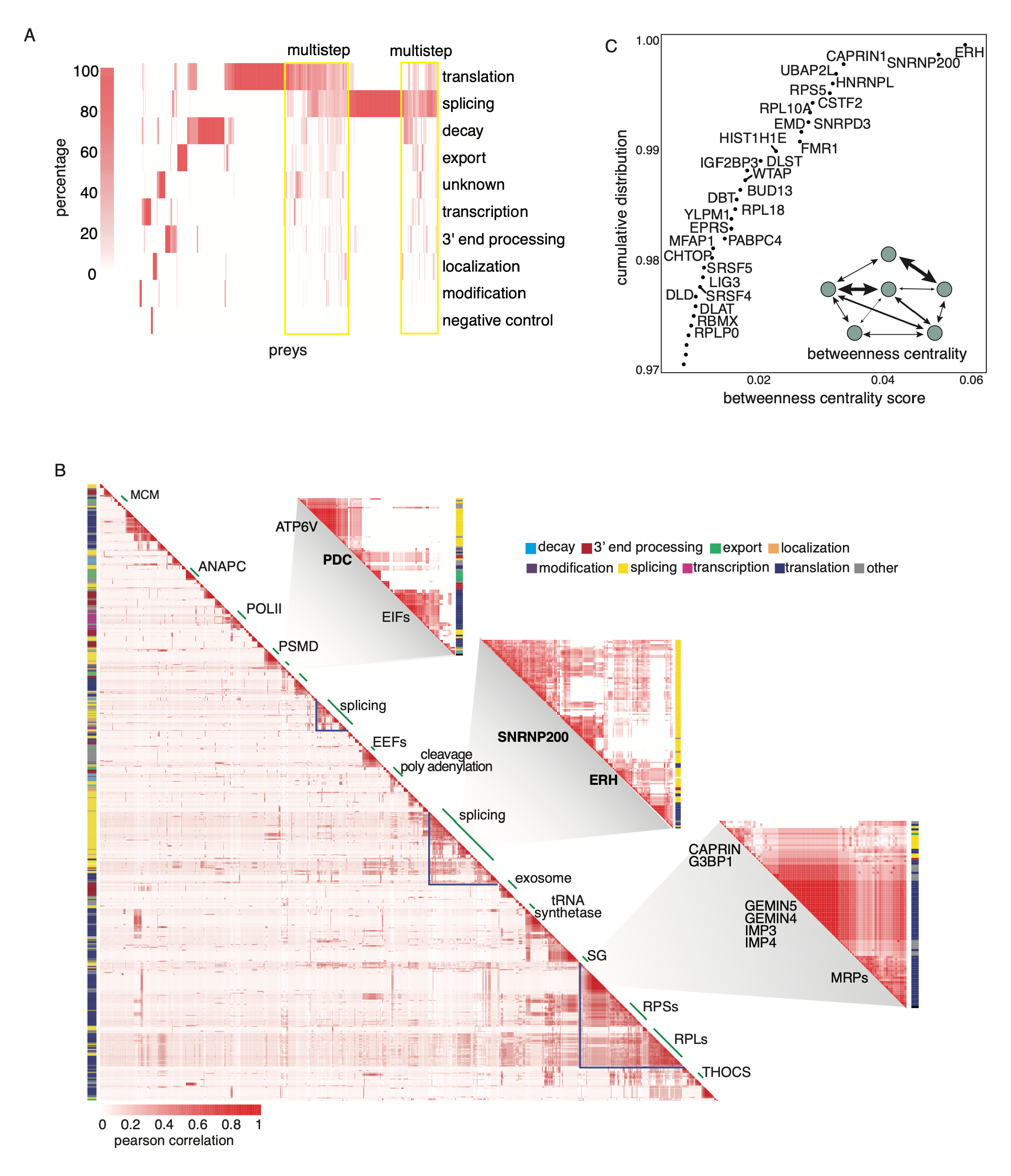
Prey-Prey correlation defines the organization of protein complexes. A. Heatmap of hierarchical clustering of the distribution of life-cycle steps associated with each prey based on the life-cycle associations of its interacting bait(s). B. Heatmap showing the Pearson correlation of all the preys’ interactions from the entire direct, RNA-mediated, and −shielded network. Indicated clusters are well-known CORUM complexes. Enlarged view of off-diagonal cluster showing preys whose interactions correlated to pyruvate dehydrogenase complex (PDC), SNRNP200, and ERH. C. Cumulative distribution plot of the betweenness centrality scores. The top 3% of RBPs with the highest scores are plotted.

Given the relatively small number of baits compared to preys in our PPI network, we next adopted a prey-centric approach to further explore the relationships between preys. We correlated all pairwise interactions (including direct, RNA-mediated, −shielded, and undetermined interactions) between each bait and prey using the Pearson correlation coefficient (Supplemental Table 6). Our rationale was that preys interacting with similar baits likely form functional RNP complexes. The heatmap of the correlation of all preys (n = 1103) produces dense clusters corresponding to well-defined protein complexes that function across the mRNA life-cycle (Figure 4B), such as the anaphase-promoting complex (ANAPAC), eukaryotic initiation factors (EIFs), and the small and large ribosomal subunits. Each prey is marked with a color representing the mRNA life-cycle step to which it was assigned using the previously mentioned unsupervised clustering. Some preys do not distinctly divide into mRNA life-cycle steps, suggesting that they might be multifunctional RBPs. Non-uniform clusters with many off-diagonal interactions were also observed, implicating that these RBPs or RNPs are involved in multiple functions and complexes. One of the off-diagonal clusters connects the pyruvate dehydrogenase complex (PDC) with many EIF proteins, suggesting an uncharacterized role for the enzymatic complex in mRNA translation. We also observe that stress granule proteins, such as G3BP1, FMR1, and CAPRIN, interact with ribosomal subunits and many other RBPs that regulate translation (Figure 4B). Taken together, correlating preys into an interaction map not only recapitulates well-known complexes, but more importantly, highlights the multifunctional nature of multiple RBPs.

### Betweenness centrality analysis reveals RBPs that function across multiple mRNA life-cycle steps

To programmatically identify RBPs that function across multiple steps in the mRNA life-cycle, we computed a betweenness centrality score (BCS) on each node (baits and preys) in our network. The BCS is calculated by finding all connected pairs of proteins between different life-cycle steps and then, for a given protein, measuring the number of the shortest paths from these connected pairs that pass through the given protein (see methods). The scores were normalized based on the "traffic load" (e.g., number of interactions) between the life-cycle steps (Figure 4C and Supplemental Figure 3B). We hypothesize that RBPs with high scores are multi-functional proteins that “bridge” RNA life-cycle steps and that the loss of intermediates will significantly impact the network and subsequent RNA processing events. The RBPs with the highest centrality scores included ERH, SNRNP200, HNRNPL, CAPRIN1, and UBAP2L (Figure 4C and Supplemental Table 7). To support our claim that proteins with high BCS values are associated with multiple mRNA life-cycle steps, we conducted an analysis on the bait life-cycle step distribution for the top 30 proteins with the highest centrality scores (Supplemental Figure 3C). This analysis demonstrates that a significant majority of these proteins are associated with baits across multiple life-cycle steps, providing strong evidence that these proteins exhibit a high degree of multifunctionality and may serve as potential bridges between multiple steps.

### Enhancer of rudimentary homology (ERH) scaffolds nuclear speckle formation necessary for RNA splicing and nuclear export

We next performed a more in-depth analysis of the roles played by some of these RBPs with high BCS values. Our PPI network indicates that the protein ERH interacts with multiple proteins across mRNA life-cycle steps, including transcription, splicing, degradation, and nuclear export. Specifically, ERH interacts directly with polymerase II (POLR2A); the splicing factors SF1, SNRNPC, SART1, MFAP1; components of post-splicing complexes and mediators of nonsense-mediated decay CASC3 and UPF2; microRNA processor DGCR8; as well as mRNA export and TREX component CHTOP, POLDIP3, and ALYREF (Figure 5A and Supplemental Figure 4A). These findings are consistent with previous reports that suggest human ERH has multiple protein partners (Graille 2022; Pang et al. 2022; Fang and Bartel 2020; Kwon et al. 2020; Kavanaugh et al. 2015). ERH is one of the most conserved proteins in metazoans, with one amino acid difference between humans and zebrafish (Wojcik et al. 1994), and it forms a butterfly-like homodimer with many protein-binding sites (Graille 2022; Hazra et al. 2020). ERH has also been previously described to form a complex with Drosha and DGCR8 and shown to be important for processing suboptimal miRNAs (Kwon et al. 2020). Depletion of ERH was previously shown to affect differential splicing of thousands of transcripts and result in an increase in the expression of γH2AX, a marker for DNA damage (Kavanaugh et al. 2015). However, ERH does not have a canonical RBD.

**Figure 5.**
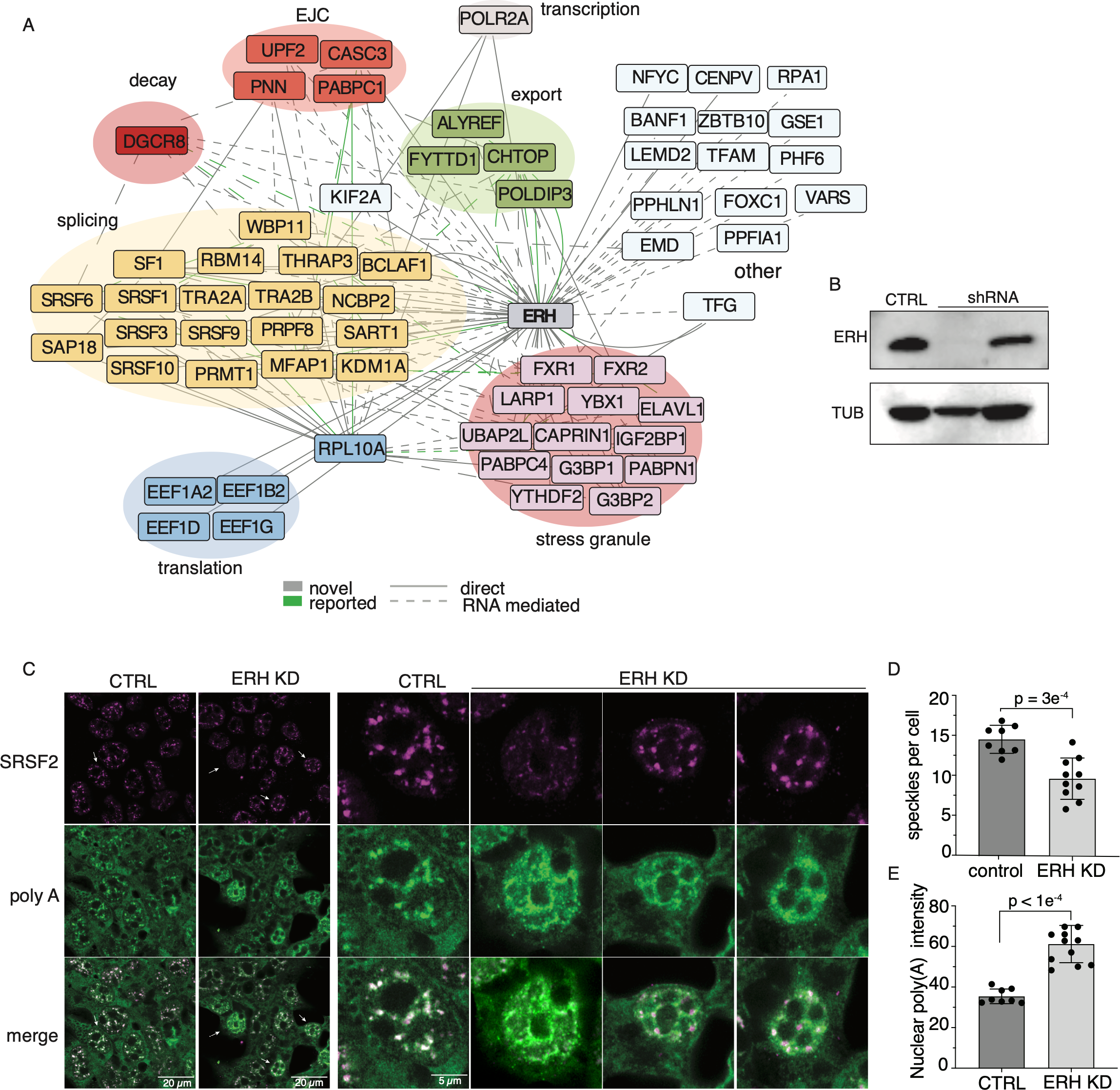
ERH interacts with many RBPs to mediate nuclear speckle organization and mRNA export. A. Cytoscape plot of ERH direct and RNA-mediated interactions. Solid lines denote direct interactions, dotted lines represent RNA-mediated interactions. Proteins are colored by life-cycle steps or functional complexes. B. Immunoblot of ERH in HEK293XT showing the shRNA knockdown vs control. C. Immunofluorescence of nuclear speckle marker SRSF2 (SC35) and poly(A) RNA in HEK293XT. D. Top panel: Quantification of nuclear poly(A) signal intensity in knockdown cells compared to control. Bottom panel: Quantification of number of nuclear speckles in control cells vs ERH knockdown cells, p-value < 3e^-4^ (unpaired t test).

To evaluate if ERH itself directly binds RNA, we performed Crosslinking and Solid Phase Purification (CLASP) in HEK293XT cells to capture proteins that have been covalently crosslinked (UV 254 nm) to RNA after high denaturing washes (Kim et al. 2021). Compared to our positive controls (PCF11 and YTHDC1), ERH does not bind a significant amount of RNA (Supplemental Figure 4B). This was further confirmed by our attempt at ERH eCLIP, where we found very few enriched targets, with the only enriched RNA target being the highly expressed, nuclear speckle, non-coding RNA *7SK*. Taken together, our findings suggest that ERH is not a significant RBP and likely functions as a scaffold protein that facilitates RBP interactions.

Our identification of ERH interacting with TREX components supports the previous finding and cryo-EM structure showing that recombinant TREX-mRNA complex co-purifies with ERH (Pacheco-Fiallos et al. 2023). TREX is highly conserved and an essential player in mRNP biogenesis, regulating and interacting with components of mRNA processing steps from 5’ cap binding to mRNA export from the nucleus. TREX components act as a platform within or on the periphery of nuclear speckles, facilitating splicing and mRNA export (Dias et al. 2010; Heath, Viphakone, and Wilson 2016; Galganski, Urbanek, and Krzyzosiak 2017; Zuckerman et al. 2020). Clustering and annotating the prey-prey correlation of the interacting partners of ERH, we see over 30 proteins known to be involved in nuclear speckle formation (Supplemental Figure 4C). However, ERH’s role as a TREX component and how it affects nuclear speckles is poorly understood (Galganski, Urbanek, and Krzyzosiak 2017).

We thus measured if the absence of ERH affects nuclear speckle formation. Nuclear speckles are membraneless organelles that undergo liquid-liquid phase separation (LLPS) and are highly enriched with splicing factors (Strom and Brangwynne 2019). Depletion of ERH protein using shRNAs resulted in an 85% reduction in ERH protein levels (Figure 5B). Nuclear speckles were visualized using an antibody against nuclear speckle core protein SRSF2 (also known as SC35) (Yamamoto et al. 2016). In the absence of ERH, the number of nuclear speckles was significantly reduced, with rounding and enlargement of speckles (Figure 5C and 5D) (Shav-Tal et al. 2001; Spector and Lamond 2011; Lamond and Spector 2003; Spector, Fu, and Maniatis 1991; Ferreira, Carmo-Fonseca, and Lamond 1994; Melčák et al. 2000).

Since nuclear speckles and TREX components are known to facilitate mRNA export out of the nucleus, we investigated the localization of poly(A) RNAs into the cytoplasm using fluorescent oligo-dT probes (T30-ATT-488). In the control cells, the poly(A) signal within the nucleus considerably overlaps with the SRSF2 signal, supporting mRNA accumulation in these nuclear compartments. Knockdown of ERH significantly increased the amount of nuclear poly(A) RNA in the nucleus (Figure 5E), suggesting a reduction in the nuclear export of mRNAs. We conclude that one of ERH’s unexpected major roles is to mediate nuclear speckle formation and facilitate the export of poly(A) RNA out of the nucleus.

### The canonical core spliceosome protein SNRNP200 interacts with RNA granule proteins CAPRIN and G3BP1

The SNRNP200 protein exhibits the second highest BCS and is part of the U5 snRNP that unwinds the U4/U6 RNA duplex (Laggerbauer, Achsel, and Lührmann 1998; Raghunathan and Guthrie 1998). Numerous structural studies show that SNRNP200 and PRPF8 form a complex that stimulates SNRNP200 unwinding activity (Malinová et al. 2017; Preussner et al. 2022; Bergfort, Hilal, et al. 2022; Bergfort, Preußner, et al. 2022). Our network indicates that SNRNP200 directly interacts with U5 snRNP proteins PRPF8 and CD2BP2. SNRNP200 also interacts in an RNA-shielded manner with the splicing factors EFTUD2, AQR, and SF3B4 (Figure 6A, Supplemental Figure 5A), which are part of the splicing B complex (Bergfort, Preußner, et al. 2022). Interestingly, our network also showed that SNRNP200 interacts directly with proteins participating in RNA granule assemblies, such as CAPRIN1, G3BP1, USP10, and FMR1, and in an RNA-mediated manner with IGF2BP1, IGF2BP3, G3BP2, ATXN2L, FXR1, and FXR2. These interactions concur with the recent finding that SNRNP200 interacts with the central SG protein CARPIN1 in neuronal-like cells (Vu et al. 2021), especially during oxidative stress conditions. Since G3BP1 and CAPRIN are mostly found in the cytoplasm, our results suggest that SNRNP200 comprises two distinct complexes, one in the nucleus performing its well-known function in the U5 snRNP, and one in the cytoplasm whose role has not yet been explored.

**Figure 6.**
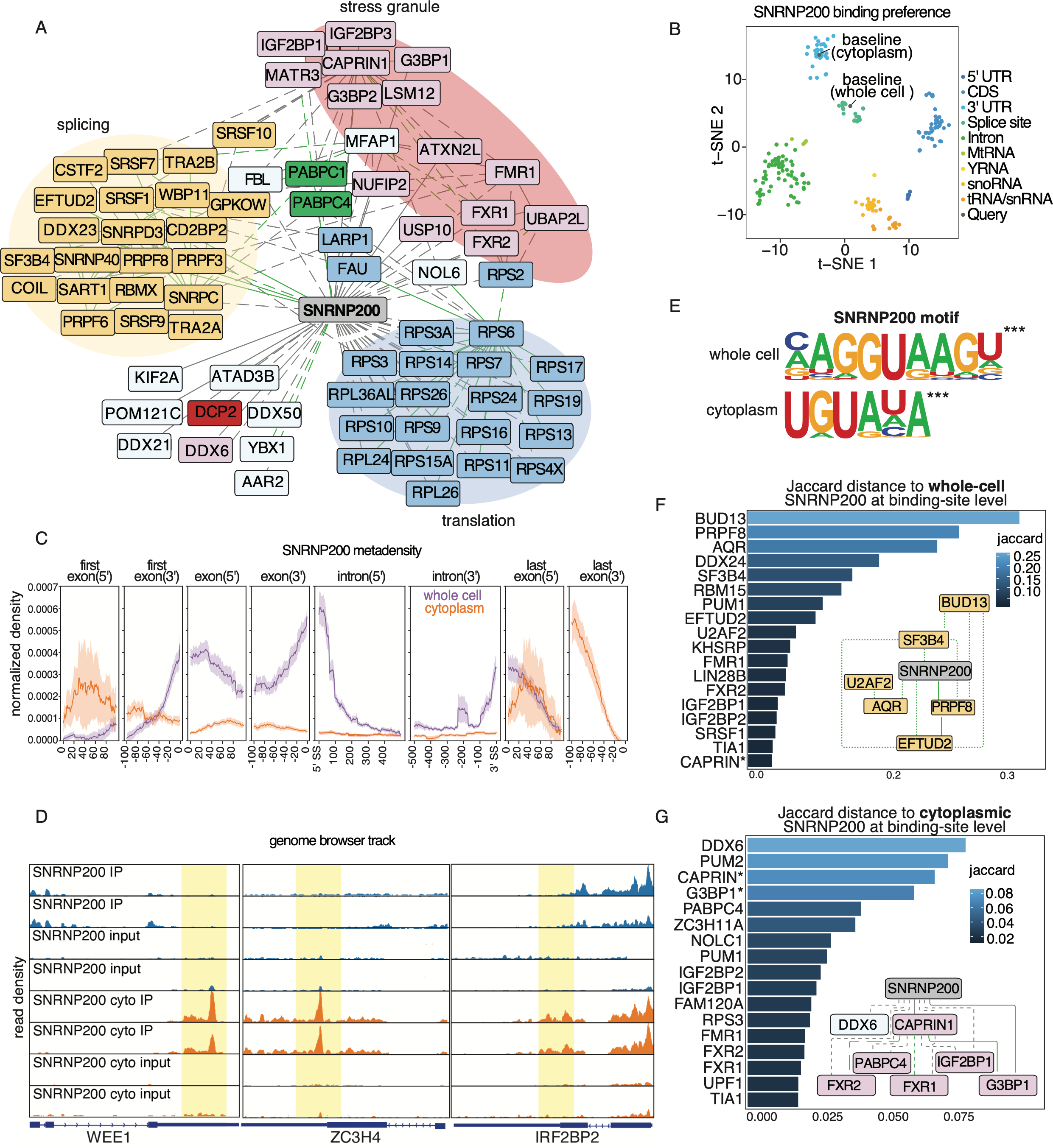
Cytoplasmic SNRNP200 has distinct binding profiles and protein partners. A. Cytoscape plot of SNRNP200 network showing only direct and RNA-mediated interactions for clarity. Solid lines denote direct interactions, dotted lines represent RNA-mediated interactions. Proteins are colored by their previously reported mRNA lifecycle step. B. tSNE plot of SNRNP200 whole-cell and cytoplasmic SNRNP200 eCLIP samples in the context of published ENCODE eCLIP datasets. C. Metadensity profiles of whole-cell (purple) vs cytoplasmic SNRNP200 (orange) from eCLIP data. D. Genome browser tracks of three representative transcripts (WEE1, ZC3H4, and IRFBP2) to show the eCLIP binding pattern for whole-cell SNRNP200 and cytoplasmic SNRNP200. E. Most significant (p-value) HOMER motif for whole-cell and cytoplasmic SNRNP200. F. Bargraph of the RBPs with the highest Jaccard index (e.g. most similar binding sites) to whole-cell SNRNP200. Inset shows the Cytoscape plot and interaction types of the RBPs with the highest Jaccard indices. G. Bargraph of the RBPs with the highest Jaccard index (e.g. most similar binding sites) to cytoplasmic SNRNP200. Inset shows the Cytoscape plot and interaction types of the RBPs with highest Jaccard indices.

To evaluate SNRNP200 protein-RNA target interactions separately in the nucleus and cytoplasm, we performed eCLIP assays in whole-cell and cytoplasmic fractions of HEK293XT cells. Global visualization of SNRNP200 gene region preferences, compared with all other ENCODE eCLIP datasets (∼300 RBPs), clearly shows that whole-cell SNRNP200 RNA target preferences are grouped with other proximal splice-site binding RBPs, while cytoplasmic SNRNP200 data clusters with RBPs that bind 3’UTRs (Figure 6B). Previous literature and biochemical assays have mapped and extensively characterized SNRNP200 binding to U4 and U6 snoRNAs (Laggerbauer, Achsel, and Lührmann 1998). Our observations that SNRNP200 binds splice sites supports a possible function in the catalytic activity of spliceosomes, which has only been described in yeast (Hahn et al. 2012). We found that over 80% of whole-cell SNRNP200 binding sites are near splice sites, located directly on 5’ and 3’ splice sites, or within the CDS near exon-intron boundaries (Figure 6C, Supplemental Figure 5B). Moreover, whole-cell SNRNP200 binds to most of the splice sites of the expressed transcripts (n = 8,000) within the cell. These findings suggest a crucial role for SNRNP200 in the catalytic activity of the spliceosome on most transcripts.

In contrast, we found that around 60% of cytoplasmic SNRNP200 binding sites are located within the 3’UTR (Supplemental Figure 5B), with the strongest enrichment observed around 100 nucleotides (nts) before the poly(A) tail and 50 nts after the stop codon (Figure 6C). We also generated genome browser tracks of three representative transcripts (WEE1, ZC3H4, and IRFBP2) to show the eCLIP binding patterns for whole-cell SNRNP200 and cytoplasmic SNRNP200 (Figure 6D). The top terms in the GO analysis (based on FDR) of the transcripts bound by cytoplasmic SNRNP200 are ‘viral processing’ and ‘negative regulation of RNA metabolic processing’ (Supplemental Figure 5C), suggesting that SNRNP200 regulates transcripts that encode for proteins important for RNA processing. The most significant motif for whole-cell SNRNP200 is the U1 snRNP (and 5’ splice site) binding signal NAGGUAAGN (IUPAC-IUB motif), with a p-value < 1e^-4^ (Figure 6E). The top motif for cytoplasmic SNRNP200 is interestingly the Pumilio response element (PRE) UGUAHA (IUPAC-IUB motif). These results indicate an increase in signal in the 3’UTR for cytoplasmic SNRNP200, while most of the signal for whole-cell SNRNP200 is concentrated near the 5’ splice sites. Our results also highlight the importance of performing eCLIP in different subcellular locations so as to capture differences in how an RBP binds RNA depending on subcellular context.

As RBPs that interact are more likely to bind to similar transcripts, we calculated the Jaccard index on binding sites bound by either whole-cell or cytoplasmic SNRNP200 (Figure 6F and G) and compared them with the K562 ENCODE eCLIP datasets. Whole-cell SNRNP200 shared the highest number of binding sites with splicing factors and B complex components BUD13, PRPF8, and AQR, which we see in our PPIs (Schmitzová et al. 2023). Conversely, cytoplasmic SNRNP200 displayed the strongest correlation with the P-body proteins DDX6 and PUM2 (Hubstenberger et al. 2017), as observed from ENCODE data, and the RNA granule proteins CAPRIN and G3BP1, for which we generated the CLIP data. These findings suggest that depending on its subcellular localization, SNRNP200 demonstrates target occupancy differences in both the maturation of RNA transcripts and the location of binding sites. Excitingly, these observations support the concept that SNRNP200 forms two distinct RNP complexes, each with unique functions.

To explore the transcriptome-wide binding profiles of these interacting RBPs, we plotted Metadensity profiles of transcripts that were bound by SNRNP200 and G3BP1, SNRNP200 and CAPRIN, or SNRNP200 alone (Supplemental Figure 5D) (Her, Boyle, and Yeo 2022). This analysis tests the hypothesis of whether specific partner proteins influence the binding profile or binding enrichment level. On transcripts that both G3BP1 and SNRNP200 bind, G3BP1 binds the 5’ end of the last exon within the coding region (Figure S6D), and SNRNP200 is highly enriched near the 3’ end of the last exon (3’UTR) compared to transcripts that SNRNP200 binds without G3BP1. CAPRIN also has a more pronounced binding profile in the 5’ and 3’ ends of the last exon and within the first exon. Altogether, these data show that transcripts that are bound by both G3BP1 and SNRNP200 or CAPRIN and SNRNP200 are more highly enriched than transcripts that were solely bound by either of the RBPs.

### SNRNP200 localizes to stress granules independent of its spliceosome partners

Given that SNRNP200 interacts with multiple proteins found within stress granules, we wanted to confirm these interactions using immunofluorescence (IF) and high-resolution microscopy. We show that while SNRNP200 exhibits both nuclear and cytoplasmic presence, it co-localizes with the predominantly cytoplasmic stress granule protein G3BP1 in unstressed HEK293XT cells. However, the co-localization is even more pronounced under arsenic stress conditions (Figure 7A). These findings extend beyond HEK293XT cells, as a previous study observed that SNRNP200, G3BP1, and CAPRIN also co-localize in the neuronal-like cell line SH-SY5Y (Vu et al. 2021).

**Figure 7.**
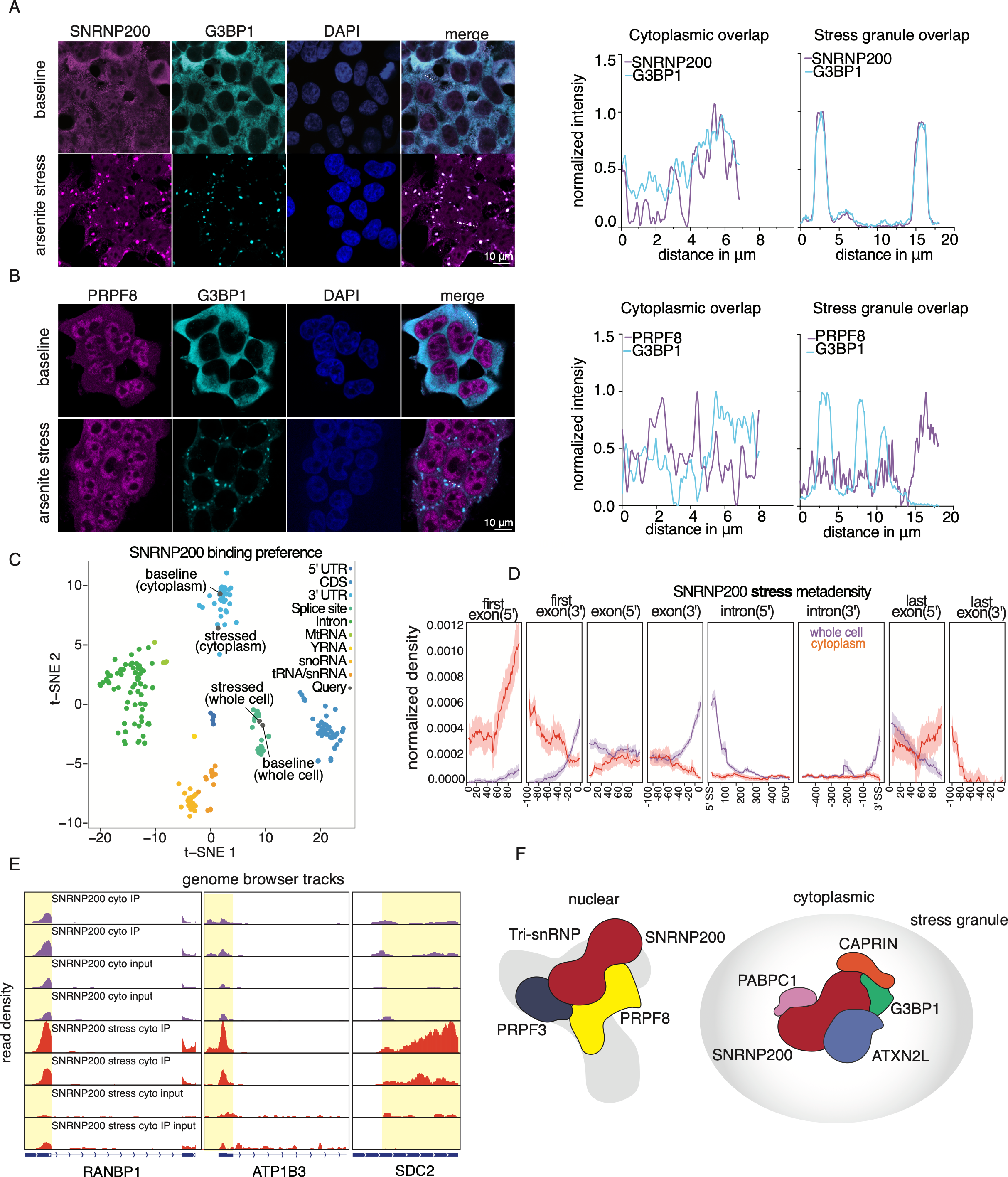
Stress granule factor SNRNP200 binds the 5’UTRs and 3’UTRs of targets during arsenic stress. A. Immunofluorescence of SNRNP200, G3BP1, poly(A) RNA, and DAPI in HEK293XT at baseline and 1 hour after arsenic stress. Line scan (depicted) analysis indicates levels of localization in µm. B. Immunofluorescence of PRPF8, G3BP1, poly(A) RNA, and DAPI in HEK293XT at baseline and 1 hour after arsenic stress. Line scan (depicted) analysis indicates levels of localization in µm. C. tSNE plot of SNRNP200 whole-cell and cytoplasmic SNRNP200 eCLIP samples during arsenic stress in the context of published ENCODE eCLIP datasets. D. Metadensity profiles of whole-cell SNRNP200 (purple) vs cytoplasmic SNRNP200 (red) during stress. E. Genome browser tracks showing eCLIP read density for cytoplasmic SNRNP200 +/- stress at the transcripts RANBP1, ATP1B3, and SDC2. F. Schematic of the two RNP complexes, from the nucleus and cytoplasm, in which SNRNP200 is a member.

Moreover, we confirmed that SNRNP200 depletion affects stress granule formation, consistent with previous findings (Wheeler et al. 2020) (Supplemental Figure 6A). SNRNP200 tightly interacts with the splicing factor PRPF8, and these two proteins have been shown to assemble in the cytoplasm and be transported to the nucleus (Malinová et al. 2017). However, our PPI network and IF data (Figure 7B) show that PRPF8 does not localize with granule proteins at baseline or during stress. Together with our co-IP data from our network, these observations suggest that G3BP1 and SNRNP200 form a functional RNP in the cytoplasm without PRPF8 in what is likely a distinct role outside of the spliceosome.

### SNRNP200 binds a subset of early response transcripts during stress

Next, we identified the RNA targets of SNRNP200 in HEK293XT cells during one hour of arsenic stress. We mapped the binding sites of both whole-cell and cytoplasmic SNRNP200 and compared them to the targets from unstressed HEK293XT cells (Supplemental Figure 6B and C). Whole-cell SNRNP200 has a very similar profile in both stress and non-stress, enriching for both 5’ and 3’ splice sites and binding a similar number of transcripts between conditions (Figure 7C and Supplemental Figure 6C). The most significant binding motif from stressed whole-cell SNRNP200 is the same as it was at baseline (Supplemental Figure 6D). However, during arsenic stress, cytoplasmic SNRNP200 binds transcripts in the 3’UTR and the 5’UTR in a manner distinct from the binding pattern of cytoplasmic SNRNP200 at baseline (Figure 7C). Specifically, the Metadensity plot shows enrichment of binding 80 nts into the first exons (Figure 7D).

Directly comparing the cytoplasmic RNA targets during baseline and stress, we see a distinct set of mRNAs that are bound in just the stress condition, with nearly 60 transcripts that gain binding sites during arsenic stress. Genome browser tracks of three representative transcripts (RANBP1, ATP1B3, and SDC2) show the cytoplasmic SNRNP200 binding pattern at baseline and stress (Figure 7E). GO analysis of these transcripts identified the most significant terms as ‘protein localization to endoplasmic reticulum’ and ‘viral processing’ indicating that SNRNP200 might play a role in protein trafficking and RNA metabolism (Supplemental Figure 6E and F). Altogether, these observations emphasize the unexpected and multifunctional roles that SNRNP200 plays across the mRNA life-cycle (Figure 7F).

## DISCUSSION

In this study, we developed an RNA-aware PPI network centered around 92 RBPs involved in several distinct stages of the mRNA life-cycle. RNase treatments allowed us to map the RNA-dependency of these PPIs. We obtained a higher percentage of RNA-mediated interactions from RNA-scaffolded complexes such as the spliceosome and ribosome, compared to the primarily direct interactions found in stable complexes such as the exosome and RNA pol II complex. An unexpected third category of interactions were only reliably detected upon RNase treatment. Possible explanations for these RNA-shielded interactions include 1) epitope exposure in the bait’s non-RNA bound state, 2) liberation of the bait from inaccessible large RNP complexes upon partial RNA digestion, or 3) spurious post-lysis interactions driven by RNase treatment. Many of these RNA-shielded interactions are part of large RNP complexes, such as the spliceosome and ribosome, and are previously reported in published PPI networks, indicating that they represent biologically functional interactions detected due to bait and epitope unmasking. However, we note that we cannot rule out artifactual post-lysis interactions for this or the other two interaction types.

We augmented our interactome with SEC-MS data that enabled us to add substructure to the IP-MS scaffold by the addition of prey-prey interactions. SEC-MS allowed us to provide orthogonal support for interactions within the IP-MS network. Furthermore, as we have added prey-prey resolution, we were able to gain a fuller picture of the prey proteins’ interactions. For example, if a prey is associated with multiple baits, its interactions with other preys can indicate if it functions within multiple RNPs or if it forms more transient interactions with other baits outside of its primary function. This IP-MS/SEC-MS combination allows us to predict RNA-binding functions for the large number of our PPI preys that have been detected as RBPs by previous RNA-interactome studies but not yet functionally characterized.

Our PPI network is reliable by a variety of parameters. We show significant levels of recovery of previously reported interactions from a range of databases and publications, and find significant levels of subcellular co-localization for interacting proteins in our interactome. When we asked how our interactome overlaps with complementary methodology, such as ENCODE eCLIP datasets, we found that measured interactors showed more significant co-binding to mRNA targets. Notably, the direct interactors displayed the highest overlap with mRNA targets in the ENCODE dataset. Finally, we found that most of the preys in the network are predicted to have RNA binding function. Taken together, these analyses gave us confidence in our network.

Global evaluation of network properties revealed general principles about mRNA life-cycle PPI behavior. Based on annotations in the Human Protein Atlas (HPA) (Thul et al. 2017), we found that proteins that interact in our network are significantly more co-localized than expected by chance, and that direct interactions had the largest co-localization frequency. We did note however, that as also observed in the OpenCell network, which mapped protein localizations in HEK293 cells (Cho et al. 2022), many interactions did not have any localization overlap. While lysis artifacts likely contribute to this lower co-localization, these results do indicate that current localization information may not fully capture the diverse range of subcellular localizations that proteins can have within a cell. Factors such as cell-type differences, RBPs with multiple functions, or variations in protein stoichiometry across different subcellular locations may contribute to this observation. For instance, we found that SNRNP200, which we observed to co-localize with different proteins in the cytoplasm and nucleus, is only assigned a nucleoplasm localization in the available dataset (HPA). This example supports the hypothesis that many RBPs may have multiple simultaneous or context-dependent subcellular localizations to carry out functions at different stages of the mRNA life-cycle. These findings highlight the complexity of protein localization and the need for further investigation to better understand the dynamic and multifaceted nature of RBPs in cellular processes.

Preys in our network were strongly enriched for RBPs as predicted by available databases and classifiers. Clustering these preys agnostic to the bait-assigned life-cycle stages revealed functional hubs representing known complexes within mRNA life-cycle stages. We learned that about 50% of preys are connected to at least two life-cycle stages, indicating a greater degree of crosstalk between the different stages and processes of the mRNA life-cycle than has yet been appreciated. Based on the high levels of connectivity we asked if the stoichiometric ratios of interacting protein abundances reflected the varying levels of connectivity we saw within the network. We found that core complexes had stoichiometries at ratios close to 1:1 while our most connected proteins (those that bridge the interactome and life-cycle stages; Figure 4B) had substoichiometric ratios. This is in line with previous work that has also shown that substoichiometric interactions define the overall structure of PPI networks (Hein et al. 2015). Interestingly, when we looked at the different types of interactions, we found that direct interactions had ratios close to 1:1, similar to core complexes, while RNA-mediated interactions had ratios below 1:1 and perhaps play a greater role in connecting the mRNA life-cycle. Follow-up work on ERH and SNRNP200, the two most connected proteins in the network, showed that these low-stoichiometry PPIs play unexpected and crucial roles in multiple mRNA life-cycle stages.

We demonstrated one such low-stoichiometry interactor, ERH, interacts with multiple RBPs involved in various stages of mRNA biogenesis. By both CLASP and eCLIP analysis, we confirmed that ERH’s function is primarily mediated through its interaction with proteins and other RBPs rather than ERH directly binding to RNA. We showed that depletion of ERH led to a significant reduction in nuclear speckles and impaired nuclear export of mRNAs. These findings indicate that ERH plays a crucial role in facilitating the export of mRNA molecules from the nucleus. Our network analysis also supports the recent discovery of endogenous tagged DGCR8’s interaction with ERH. Interestingly, knockdown of ERH resulted in decreased levels of mature miRNAs while having no effect on pri-miRNA levels (Kwon et al. 2020). The diverse functions of ERH can be attributed to its structural characteristics and its ability to homodimerize (Hazra et al. 2020), which likely allows it to bind to numerous protein partners that ultimately dictates its specific role.

We have uncovered unexpected and exciting insights into the intricate interactions and multifaceted functions of SNRNP200 in the cell. While previous research has focused on SNRNP200’s role in binding and unwinding the U4-U6 RNA duplex, our eCLIP assays and PPI network revealed that it also binds to the splice sites of most transcripts, suggesting a possible involvement in the catalytic activity of the spliceosome. Our transcriptome-wide results in mammalian cells amplify findings by Hahn et al. that demonstrated that the yeast homolog of SNRNP200, Brr2, crosslinks to splice sites and that its mutation impairs the second transesterification step of introns (Hahn et al. 2012). Our study also characterized the cytoplasmic function of SNRNP200, as evidenced by its interactions with RNA granule proteins at cellular baseline and its binding to the 3’UTRs of various transcripts in the cytoplasmic fraction. Under cellular stress, SNRNP200 was found to enrich for 5’UTRs of a specific set of mRNAs. Interestingly, SNRNP200 has been found enriched in amyotrophic lateral sclerosis patient neural protein aggregates (Vu et al. 2021; Lachén-Montes et al. 2020), which places it at the intersection of toxic protein aggregation and mRNA processing dysregulation.

Although our interactome represents a significant advancement in understanding mRNA processing and regulation, there are important considerations that warrant further investigation. Firstly, while we try to minimize artifacts in our interactome by using endogenous antibodies and provide orthogonal support from two different methods - IP-MS and SEC-MS - we cannot rule out that some of the interactions may be technical artifacts due to cellular lysis, which affects both IP-MS and SEC, or due to antibody cross-reactivity artifacts in IP-MS. Another area requiring further exploration is the characterization of RNA-shielded interactions. Understanding the nature of these interactions and their biophysical properties will require additional research.

In summary, we have developed a novel RNA-aware protein-protein interactome centered around RBPs and focused on the mRNA life-cycle. Furthermore, our network analysis has led to the identification of key RBPs and their associated proteins, such as ERH and SNRNP200, while generating functional hypotheses for uncharacterized proteins. Our approach, combining two complementary methods, not only provides insights into fundamental principles governing mRNA processing and regulation, but also serves as a valuable resource for the scientific community.

## AUTHOR CONTRIBUTIONS

KB, GWY, MJ conceived the study. KRothamel, LS, WJ, BB, and AA developed the software, tested the software, collected data, analyzed data and visualized results. KB, LS, and KRothamel designed wet-lab validation experiments. KRothamel, LS, KB, YM, JKK, AM, KRhine, NA, KD, EW, MM, JS, and EDM carried out the experimental work. KRothamel, LS, and WJ designed computational validation experiments. KRothamel, LS, and WJ carried out the computational validation experiments. LS and KRothamel wrote the original manuscript draft. GWY, MJ, KB reviewed and edited the manuscript. GWY and MJ acquired the funding and supervised the study.

## ACKNOWLEDGMENTS

G.W.Y. is supported by NIH R01 HG004659, U24 HG009889 and an Allen Distinguished Investigator Award, a Paul G. Allen Frontiers Group advised grant of the Paul G. Allen Foundation. M.J. is funded by the NIH (R35GM128802; R01AG071869 and R01HG012216), NSF (Award 2224211) and Columbia startup funding. K.Rothamel is supported by NIH T32CA067754. B.B. was supported by NSF-GRFP (Award DGE2036197).

## Supplemental Figure Legends

**Supplemental Figure1. The IP-MS network significantly recapitulates known PPIs across life-cycle stages and interaction types**

A. Schematic of the experimental workflow. The IP was performed in the presence and absence of RNase using endogenous antibodies followed by LC-MS/MS for each bait. IPs against the baits were performed in triplicate for both with and without RNase conditions and IgG controls were performed in enough replicates in with and without RNase conditions to meet the requirements of the TMT mixing scheme. IP samples were multiplexed into TMT-11 mixes with a spike-in channel made from a mix of all IPs in the dataset, 3 RBP IPs with and without RNase and 2 IgG controls with and without RNase. Raw data was searched using the SpectroMine software and the data was normalized. High confidence interacting proteins were identified as those preys that have normalized intensities greater than 1.645 sd in at least 2 replicates and that were significantly enriched compared to control and failed IPs (see methods for details).

B. The percentage of literature supported interactions in the IP-MS network and by interaction type for a number of different databases and previously published large-scale interactomes.

C. The distribution of different types of interactions for the baits assigned across the mRNA life-cycle (left), and the distribution of interaction types for those PPIs with literature support (right).

D. Percent of CORUM supported interactions in the network compared to randomly shuffled interactions of all the proteins in our network for n = 1000 tests, p-value < 0.001.

E. Histograms of the number of baits or unique bait life-cycle stages for the preys in the IP-MS network.

F. Prediction of RNA binding for preys in the interactome compared to proteins measured in the total HEK293XT proteome. RNA binding prediction by HydRa scores (greater scores indicate more confidence of RNA binding; score >0.89 indicates predicted RBP). K-S statistic: p-value < 2.2e^-16^.

**Supplemental Figure 2. The IP+SEC network has significant levels of co-localization and different interaction stoichiometries between complex hubs and connector proteins**

A. Schematic of the experimental workflow. Lysates from HEK293XT cells are clarified and buffer exchanged into SEC buffer before being run on a SEC column with fraction collection. Fractions are processed for mass spectrometry and run on LC-MS/MS in DIA mode. Raw data is searched using Spectronaut software and protein co-elution is evaluated by CCprofiler at an FDR of 0.1 and q-value < 0.1 against a dataset of decoy interactions.

B. Percent of interactions by interaction type that are supported by SEC-MS and IP-MS compared to those only found in the IP-MS network.

C. Percent of supported interactions in the SEC-MS network compared to randomly shuffled interactions of the proteins within our network for n=1000 tests, p-value < 0.001.

D. Percent of CORUM supported interactions in the SEC-MS network compared to randomly shuffled interactions for n=1000 tests, p-value < 0.001.

E. The percentage of literature supported interactions in the SEC-MS, IP-MS, overlap, and combined networks for a number of different databases and previously published large-scale interactomes.

F. Percent of PPIs with fully or partially co-localized annotations in the Human Protein Atlas in the IP+SEC-MS network and in the previously published large-scale Bioplex 3.0 interactome compared to randomly shuffled interactions for n=1000 tests, p-value < 0.001.

G. The stoichiometric ratios in the IP-MS data for the 5, 10, and 15 most connected proteins in the network are significantly lower (adjusted p-value Student’s t-test) than the ratios of the rest of the network. Medians are plotted in red.

H. Stoichiometric ratios of interactions found in the network. The log2 ratios of the intensities of the prey over bait in the IP-MS data are plotted by interaction type with adjusted Student’s t-test significance values. Medians plotted in red.

I. IP+SEC-MS network interactions found in CORUM.

**Supplemental Figure 3. Bait-and Prey-centric analyses link proteins to particular life-cycle steps and complexes**

A. Heatmap of hierarchical clustering grouping of the distribution of life-cycle steps associated with each prey based on the life-cycle associations of its interacting bait(s).

B. Heatmap of correlation between bait-bait interactions.

C. Cumulative distribution plot of the traffic flow normalized betweenness centrality scores. The top 3% of RBPs are plotted.

D. Subset of Figure 3A plotting the top 30 preys with the highest centrality score. For each prey, the percent of their baits from each life-cycle step is plotted.

**Supplemental Figure 4. ERH, a non-RBP, associates with nuclear speckles proteins**

A. Cytoscape plot of all the ERH interactions (including RNA shielded, mediated and direct interactions). Groups of RBPs are colored by GO ontology associated with mRNA life-cycle steps.

B. Prey-prey plot of the proteins that interact with ERH. Each protein is colored based on the hierarchical clustering assignment to a particular life-cycle step. Proteins are also colored if they were found to be in nuclear speckles (green).

C. Top: Total protein blot of CLASP experiment. Bottom: Immunoblot of the CLASP experiment staining for the RBPs YTHDC1, PCF11 and the negative control protein TUB4A and experimental protein ERH.

**Supplemental Figure 5. SNRNP200 interacts and co-binds transcripts with stress granule proteins CAPRIN and G3BP1**

A. Full cytoscape network of SNRNP200 interactions. Proteins are grouped based on GO ontology terms.

B. Barplot of the percentage of SNRNP200 windows that mapped to specific transcript features (5’UTR, CDS, 3’UTR etc.) in the whole-cell eCLIP vs the cytoplasmic eCLIP.

C. GO terms for the transcripts bound by SNRNP200 in the cytoplasmic fraction. Terms are ordered by FDR values.

D. Metadensity plots showing the binding profiles of G3BP1 (blue) or CAPRIN (green) and SNRNP200 (orange) for transcripts that are known to be bound by both RBPs or exclusively by cytoplasmic SNRNP200.

**Supplemental Figure 6. SNRNP200 binds a distinct set of transcripts during stress**

A. Immunofluroscence images counting the number of stress granules (G3BP1) in each cell in shRNA SNRNP200 knockdown and control. Bargraph of the quantification of the number of stress granules per cell across 10 images. Student’s paired t test was used to calculate if there was a difference in the means.

B. Barplot of the percentage of windows that mapped to transcript features in stressed whole-cell versus stressed cytoplasmic SNRNP200 eCLIP.

C. Upset plot of significant (p-value < 0.001 and enrichment > 3) windows from SNRNP200 eCLIP samples from whole-cell and cytoplasmic SNRNP200 at baseline and stressed conditions.

D. Most significant HOMER motif analysis of stressed whole-cell and cytoplasmic SNRNP200.

E. GO analysis for transcripts bound by cytoplasmic SNRNP200 during stress.

F. GO analysis for transcripts that are distinctly bound by SNRNP200 in the cytoplasm during stress.

## METHODS

### Cell Culture

HEK293XT (Takara Bio Lenti-X 293T, #632180) cells were cultured in DMEM containing L-glutamine and sodium pyruvate supplemented with 10% Fetal Bovine Serum, Penicillin (100 U/mL), and Streptomycin (100 μg/mL). Cells were grown to 90-100% confluency before passage and were harvested at passages 6-20.

For harvesting, confluent HEK293XT cells in a 10 cm plate were gently washed in cold 1X PBS and dissociated from the plate with a cell scraper. Cells were then pelleted in cold PBS at 500 g for 5 minutes. Cells were then resuspended in 1 ml of ice-cold PBS and transferred to 1.5 ml Protein Lo-Bind tubes to be pelleted again and flash-frozen in liquid nitrogen and stored at −80 °C.

### Immunoprecipitations

Cell pellets were thawed on ice and lysed in 400 µl of lysis buffer (150 mM NaCl, 50 mM Tris pH 7.5, 1% IGPAL-CA-630, 5% Glycerol, and protease and phosphatase inhibitors), and split evenly (∼200 µl) into separate tubes. Half of the lysis was treated with 5 µl of 10 mg/ml Rnase (Promega:527491) and both lystates (+/- RNase) were incubated on ice for 20 minutes. Each tube (+/- RNase conditions) was centrifuged at 4°C for 10 minutes at 14,000 g. The total protein concentration of each lysate was measured using a BCA assay to ensure that each sample had between 1-2 mgs of total protein.

100 µl of Dynabeads Protein G (Invitrogen:01200616) magnetic beads were washed 3 times in 1 ml lysis buffer and then conjugated to 10 µg of endogenous antibody. Bead-antibody conjugation was then added to the cell lysate and incubated overnight at 4°C on the rotator. The following day, samples were placed on a magnetic bead separator, the supernatant was removed, and samples were washed 2 times with wash buffer (150 mM NaCl, 50 mM Tris pH 7.5, 5% Glycerol) containing 0.05% IGPAL and 2 times with wash buffer without IGPAL. Beads were then incubated in 80 µl of on-bead buffer (2M urea, 50mM Tris (pH 7.5), 1 mM DTT, and 5 μg/mL Trypsin (Promega:487603)) for 1 hour at 25°C on a shaker (1000 rpm). After one hour, beads were placed on a magnetic bead separator and 80 µl of supernatant was transferred to the new tube. The beads were then washed twice with 60 μL of 2M urea and 50 mM Tris (pH 7.5) HPLC buffer. The supernatant from each wash was combined with the on-bead digest for a total of 200 µl per sample. Samples were then spun at 5000 g, transferred to a new tube, and stored at −80°C.

### Size Exclusion Chromatography

#### Preparation of samples for SEC

SEC samples were prepared as described in (Bludau et al. 2020). HEK293XT cells (20-30 million per replicate, n=3) were harvested at ∼80% confluence by scraping in ice cold 1X PBS with 5nM EDTA, washing and pelleting, before being flash frozen in liquid nitrogen and stored in −80°C. Pellets were lysed in ice cold lysis buffer (150 mM NaCl, 50 mM Tris pH 7.5, 1% IGPAL-CA-630, 5% Glycerol, same lysis conditions as for IP) supplemented with 50 mM NaF, 2 mM Na3VO4, 1 mM PMSF, and 1X protease inhibitor cocktail (Sigma) and incubated on ice for 30 minutes. Cell lysates were then clarified by centrifugation for 10 minutes at 10,000 g and 4°C, followed by ultracentrifugation for 20 minutes at 100,000 g and 4°C. Samples then underwent buffer exchange into SEC buffer (50 mM HEPES pH 7.5, 150 mM NaCl and 50 mM NaF) to dilute lysis detergents on a 30 kDa Amicon ultra-0.5 centrifugal filter (Sigma). Buffer exchange was done in iterative steps of no larger than 1:3 dilutions to reach a final dilution ratio of 1:50 (IP lysis buffer: SEC buffer). The lysate underwent a final clarification by 5 minutes of centrifugation at 17,000 g and 4°C. The supernatant concentration was measured by Nanodrop spectrophotometer (Thermo Scientific) and adjusted to 20 mg/mL.

#### SEC

Size exclusion was performed on an Agilent 1260 Infinity II system operated with Agilent OpenLAB ChemStation software (version C.01.09). 2 mg of cell lysate at 20 mg/mL was loaded onto a Yarra SEC-4000 column (Phenomenex 00H-4514-K0, 3 μm silica particles, 500 A pores, column dimensions: 300 × 7.8 mm) and fractionated in SEC running buffer (50 mM HEPES pH 7.5, 150 mM NaCl) at a flow rate of 0.5 mL/min. 100 μL fractions were collected between minutes 11 to 30 into 96 Well DeepWell Polypropylene Microplates (Thermo Scientific).

### Total Proteome Measurement

#### Lysis

HEK293XT cells from one 10 cm plate per replicate (n=3) were harvested at ∼80% confluence by scraping in ice cold 1X PBS, washing and pelleting, before being flash frozen in liquid nitrogen and stored in −80°C. Pellets were lysed in ice cold lysis buffer (150 mM NaCl, 50 mM Tris pH 7.5, 1% IGPAL-CA-630, 5% Glycerol, same lysis conditions as for IP) and incubated on ice for 20 minutes. Lysates were clarified by centrifugation at 14,000 g for 10 minutes at 4°C. Protein concentrations were measured by Pierce BCA protein assay kit (Thermo Scientific) and divided into 40 μg aliquots.

### LC-MS/MS

#### IP sample preparation

Spike-in samples (in 50 μL aliquots) were generated by combining equal volumes of all IP and control samples in the experiment. 50 μl of partially digested proteins were used per IP, and spike-in samples were processed identically. Disulfide bonds were reduced with 5 mM dithiothreitol (DTT) for 45 minutes at 600 rpm and 25°C. Cysteines were subsequently alkylated with 10 mM iodoacetamide (IAA) for 45 minutes in the dark at 600 rpm and 25°C. Samples were then further digested by adding 0.5 μg sequencing grade modified trypsin (Promega) for 16 hours at 600 rpm and 25°C. After digestion, samples were acidified with a final concentration of 1% formic acid. Tryptic peptides were desalted on C18 StageTips according to (Rappsilber, Mann, and Ishihama 2007) dried in a vacuum concentrator and reconstituted in 30 μl of 50 mM HEPES pH 8.5 for TMT labeling.

#### TMT labeling and multiplexing

TMT11 mixes were designed to minimize overlap between IP baits, such that baits A and B were not in the same mixes across 2 replicates and such that baits within life-cycle stages were well divided across mixes. Each TMT11 mix contained 1 spike-in (channel 126), 3 RNase treated IPs, 3 non-RNase treated IPs, 2 RNase treated IgG controls, and 2 non-RNase treated IgG controls.

Peptides were labeled by addition of a third of an aliquot of TMT11-131C label reagent (Thermo Scientific) in a final volume of 20% acetonitrile and incubated for 1 hour at 600 rpm and 25°C. Hydroxylamine at a final concentration of 0.3% was added to quench the reaction. The samples in each mix were combined, dried to at least 50% of the pooled volume in a vacuum concentrator, acidified with a final concentration of 1% formic acid, and desalted on C18 StageTips. The eluted pools were fully dried in a vacuum concentrator and reconstituted in 15 μl of 3% acetonitrile/ 0.2% formic acid for LC-MS/MS.

#### TMT LC-MS/MS

5 μL of total peptides were analyzed on a Waters M-Class UPLC using a 25 cm Thermo EASY-Spray column (2 μm, 100Å, 75 μm x 25 cm) coupled to a benchtop ThermoFisher Scientific Orbitrap Q Exactive HF mass spectrometer. Peptides were separated at a flow rate of 400 nL/min with a 190 min gradient, including sample loading and column equilibration times. Data was acquired in data-dependent mode. MS1 spectra were measured with a resolution of 120,000, an AGC target of 5e^6^ and a mass range from 300 to 1800 m/z. MS2 spectra were measured with a resolution of 60,000, an AGC target of 1e^5^ and a mass range from 200 to 2000 m/z. MS2 isolation windows of 0.8 m/z were measured with a normalized collision energy of 33 and a fixed first mass of 110 m/z.

#### SEC sample preparation

Protein concentrations of the SEC fractions were measured by Pierce BCA protein assay kit (Thermo Scientific). Equal volumes (∼80 μL) from each of the fractions containing proteins (10 - 66) were processed. Proteins were denatured by incubation in an equal volume of urea buffer (8 M urea, 75 mM NaCl, 50 mM HEPES pH 8.5, 1 mM EDTA) for 20 minutes at 600 rpm and 25°C in 96 Well DeepWell Polypropylene Microplates. Disulfide bonds were reduced with 5 mM dithiothreitol (DTT) for 45 minutes at 600 rpm and 25°C. Cysteines were subsequently alkylated with 10 mM iodoacetamide (IAA) for 45 minutes in the dark at 600 rpm and 25°C. Samples were diluted 1:3 with 50 mM HEPES pH 8.5 and then digested by a 1:50 (trypsin to protein) ratio of sequencing grade modified trypsin (Promega) for 16 hours at 600 rpm and 25°C. After digestion, samples were acidified with a final concentration of 1% formic acid. Tryptic peptides were desalted on C18 StageTips in 96 Well DeepWell Polypropylene Microplates, dried in a vacuum concentrator, and reconstituted in 3% acetonitrile/ 0.2% formic acid to a final concentration of 0.5 μg/μL.

#### SEC LC-MS/MS

5 μL of total peptides were analyzed on a Waters M-Class UPLC using a 25 cm Thermo EASY-Spray column (2 μm, 100Å, 75 μm x 25 cm) coupled to a benchtop ThermoFisher Scientific Orbitrap Q Exactive HF mass spectrometer. Peptides were separated at a flow rate of 400 nL/min with a 70 min gradient, including sample loading and column equilibration times. Data was acquired in data-independent mode. MS1 Spectra were measured with a resolution of 120,000, an AGC target of 5e^6^ and a mass range from 350 to 1650 m/z. 15 isolation windows of 87 m/z were measured at a resolution of 30,000, an AGC target of 3e^6^, normalized collision energies of 22.5, 25, 27.5, and a fixed first mass of 200 m/z.

#### Total proteome sample preparation

One 40 μg aliquot was used per replicate. The lysate volume was increased to 80 μL with 50 mM Tris pH 8. Disulfide bonds were reduced with 5 mM dithiothreitol (DTT) for 45 minutes at 600 rpm and 25°C. Cysteines were subsequently alkylated with 10 mM iodoacetamide (IAA) for 45 minutes in the dark at 600 rpm and 25°C. Samples were then digested by a 1:50 (trypsin to protein) ratio of sequencing grade modified trypsin (Promega) for 16 hours at 600 rpm and 25°C. After digestion, samples were acidified with a final concentration of 1% formic acid. Tryptic peptides were desalted on C18 StageTips, dried in a vacuum concentrator, and reconstituted in 3% acetonitrile/ 0.2% formic acid to a final concentration of 0.5 μg/μL for LC-MS/MS.

#### Total proteome LC-MS/MS

5 μL of total peptides were analyzed on a Waters M-Class UPLC using a 25 cm Thermo EASY-Spray column (2 μm, 100Å, 75 μm x 25 cm) coupled to a benchtop ThermoFisher Scientific Orbitrap Q Exactive HF mass spectrometer. Peptides were separated at a flow rate of 400 nL/min with a 160 min gradient, including sample loading and column equilibration times. Data was acquired in data-independent mode. MS1 Spectra were measured with a resolution of 120,000, an AGC target of 5e^6^ and a mass range from 350 to 1650 m/z. 46 isolation windows of 28 m/z were measured at a resolution of 30,000, an AGC target of 3e^6^, normalized collision energies of 22.5, 25, 27.5, and a fixed first mass of 200 m/z.

### IP-MS Analysis

#### Searches

Raw data were searched with SpectroMine v3.2 (Biognosys) using a UniProt database (Homo sapiens, UP000005640) under default TMT11 quantification settings but without cross-run normalization or missing value imputation. Protein group data were exported for subsequent analysis.

#### Normalization

Normalization and subsequent analysis was performed in R v3.6.3. Contaminants and immunoglobulin proteins were removed and samples’ raw intensity values were selected for normalization. iBAQ correction was performed in a TMT11 mix wise manner using “Top3MS1Quantity” values. Missing values were maintained as NAs. Parts per million (ppm) were calculated for each sample and a pseudocount of 1 was added before the data was log2 transformed. Batch normalization was performed next. Briefly, for each sample, all the other matched samples in the mix were averaged and subtracted from the sample in a protein-wise manner. Spike-in samples were normalized by subtraction of the entire rest of the mix as they contained a mix of RNase treated and untreated samples. Batch corrected samples were then z scored to allow easier between-sample comparisons. IP samples, where the bait protein itself was not enriched within the top 5% of measured proteins or where the data were too sparse for reliable analysis, were considered failed IPs and were removed from further analysis. After this filtering step, 92 bait proteins had at least 2 replicates in one RNase treatment condition and thus remained in the dataset.

#### Determining high confidence interacting proteins

High-confidence interacting proteins (HCIPs) for each of the 92 bait proteins were identified for both RNase treatment and non-treatment conditions. HCIPs were determined to be preys that were enriched within the top 5% of measured proteins in at least 2 of 3 replicates, and significantly enriched in the IP compared to background (as measured by the combined IgG control and failed IP signal [log2 ratio IP/background > 1; wilcox test p-value < 5e^-2^ or 1e^-2^ depending the bait signal]).

#### Characterizing the RNA-dependence of interactions

HCIPs identified in both the RNase treatment and non-treatment IPs for a particular bait were called “direct interactors.” HCIPs identified in only RNase non-treatment were called “RNA-mediated interactors,” and HCIPs identified in only RNase treatment were called “RNA-shielded interactors.” As 10 IP baits failed to enrich the bait entirely in either the RNase treatment or non-treatment condition, their interactors’ RNA-dependence could not be determined and they were labeled as “undetermined.” These HCIPs with characterized interaction types were then combined for all IP baits to generate the IP-MS network.

### SEC-MS Analysis

#### Searches

Raw data were searched by Spectronaut v16.0 (Biognosys)(Bruderer et al. 2015) in the directDIA method, using a human UniProt database (Homo sapiens, UP000005640) under default settings (BGS) but without cross-run normalization or missing value imputation. Peptide spectral matches (PSMs), peptides, and protein group data were exported for subsequent analysis.

#### Normalization

SEC fraction intensities were summarized with Spectronaut to the peptide quantities for each fraction. The quantities of each fraction for each peptide were averaged between the three replicates, removing NA values from the mean. A single UniProtID was selected for each peptide based on the protein group identified by Spectronaut - preference being given to UniprotIDs selected in the IP-MS network and prioritization identified by Spectronaut.

#### Identifying interactions

A reference database for CCprofiler (Heusel et al. 2019) was obtained by combining the binary interactions of CORUM and the IP-MS network, where each interaction belonged to one or more IP bait or CORUM complex. CCprofiler was run on the normalized SEC fractions, a UniProt reference, the molecular weight calibration for the SEC run, and the customized reference database. CCprofiler interactions were recovered from peaks with a q-value < 0.1 (and an FDR compared to decoys of 0.1) from the reference IP baits, where “subunits_detected” for each peak were converted to pairwise interactions for the SEC interaction network.

### Total Proteome Analysis

#### Searches

Raw data were searched by Spectronaut v16.046 (Biognosys) in the directDIA method, using a UniProt database (Homo sapiens, UP000005640) under default settings (BGS) but without cross-run normalization or missing value imputation. Protein group data were exported for subsequent analysis.

#### Normalization

Normalization and subsequent analysis was performed in R v3.6.3. Contaminants were removed and the “raw.Quantity” unnormalized intensity values were selected. A pseudocount of 1 was added and the data were log2 transformed. The three replicates were averaged and the resulting data were used for stoichiometry analysis.

### Generation of Network

#### Combination of IP-MS and SEC-MS networks and addition of interaction characteristics

The IP-MS and SEC-MS networks were combined such that any IP-MS interactions that were also identified in SEC-MS were collapsed into a single interaction with dual support. The network table was expanded to comprehensively report how the network reflects previously published data. Information about PPI support, bait and prey subcellular localization (Roux et al. 2018; Thul et al. 2017), eCLIP data (ENCODE) (Van Nostrand, Pratt, et al. 2020), HydRa RNA-binding prediction (Jin et al. 2023), phase separation propensity (Vernon et al. 2018), and disease associations (Piñero et al. 2020) was added to the interactome table.

#### Network analysis (Cytoscape)

The Cytoscape (Shannon et al. 2003) networks were generated by importing the interaction networks (IP-MS network and IP-MS+SEC-MS network). The source of interaction support or CORUM support was set to color the edges. The linetype was set by the IP RNA-aware interaction types. Network layout was done by default prefuse force directed layout without edge weights and with bundled edges. Network statistics were determined using Cytoscape default parameters.

### Interaction Stoichiometry Analysis

Stoichiometries of IP-MS interactions were calculated using un-normalized intensities as they are internally controlled within samples and to allow for minimal processing and identical processing between total proteome and IP-MS stoichiometries. IP-MS data were analyzed as described above for the total proteome data. Stoichiometries were calculated in log2 space by subtracting the bait intensity from the prey intensity.

### Resampling test for PPI networks

We firstly calculate the percentage of PPI pairs (bait-prey) that are reported by public PPI datasets (i.e., Mentha, HuRI, HIPPIE, and rec-Y2H) out of all our PPI pairs and referred to it as *Fexp*. We next randomly sampled protein either from HEK293 proteome which consists of proteins with pTPM > 0 in HEK293 cells reported by Human Protein Atlas or from all the preys we obtained in our entire experiments with all the baits. Within the IP-MS network, for each bait protein, we sampled the same number of random proteins (rand_preys) as the prey proteins found in our IP-MS results and generated a list of PPI pairs (i.e. [bait<-->rand_prey1, bait<-->rand_prey2, bait<- ->rand_prey3, …]). After combining the generated PPI pairs from all the bait proteins, we calculated the percentage the same as defined *Fexp* but in the generated PPI pair set and referred to it as *Fi*. We repeated the random sampling steps for 10,000 times and got the resampled distribution { *Fi* }, where i ranges from 1 to 10,000. The one-sided p-value of the test is calculated as the proportion of the resampled distribution where *Fi* was greater than or equal to *Fexp*. Similarly, within SEC identified interaction network between IP-MS identified preys, we used the same approach for the resampling test except for the random sampling step where the PPI pairs are randomly chosen from all potential protein-protein pairs without the restriction set above.

#### Life step bridge analysis

We defined a network topology-based score (called “bridge score”), modified from betweenness centrality score, to measure the potential of proteins to be the “bridges” between different life-cycle steps. The potential of a protein to be the “bridge” between different life-cycle steps was measured by a network topology-based score (referred to as bridge score) modified from the betweenness centrality score in graph theory. We defined the bridge score of protein *v* as the normalized sum of the fraction of all-pairs shortest paths between life steps that pass through *v*. Specifically, the bridge score for two specific life-cycle steps are defined as:

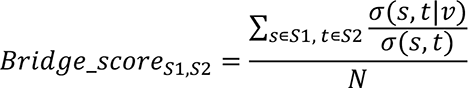

*S1* and *S2* represent the bait proteins from two given life-cycle steps. *σ*(*s*, *t*) is the number of shortest paths between node *s* and *t*. *σ*(*ν*) is the number of those paths passing through node *v.* N represents the total number of (s, t) pairs between Step 1 and Step 2.

Similarly, the bridge score measuring the protein centrality among all the life-cycle steps are defined as:

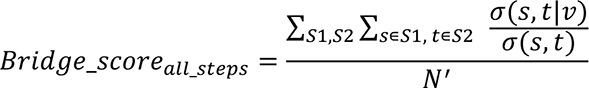

N’ represents the total number of (s, t) pairs of all pairs of steps. S1 and S2 represent the bait proteins from any two of the life-cycle steps.

To balance the traffic flows between different pair of life-cycle steps, we also defined a balanced version of the bridge score for measuring the protein centrality among all the life-cycle steps:

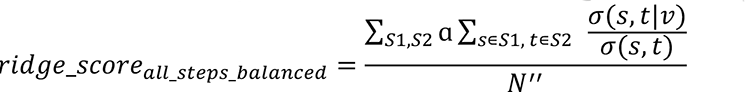

ɑ is a normalization scaling factor which is calculated as 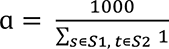, while N’’ represents the normalized total number of (s, t) pairs of all pairs of steps, which is defined as N’’ = 1000 × ∑*_S1_*,*_S2_* 1.

### eCLIP

HEK293XT cells were cultured in DMEM medium supplemented with 10% FBS at 37°C with 5% CO2. eCLIP was performed as previously described (Van Nostrand et al. 2016) with minor modifications. Briefly, cells were crosslinked with 200 mJ/cm2 UV-C light and lysed in lysis buffer (50 mM HEPES pH 7.4, 150 mM NaCl, 0.5% NP-40, 0.1% SDS, 0.5 mM DTT, 1 mM PMSF, 1x Protease Inhibitor Cocktail). Lysis was centrifuged at max speed and the supernatant was used for the immunoprecipitation with endogenous antibody conjugated to protein A/G magnetic beads. The RNA-protein complexes were washed with high salt wash buffer (50 mM HEPES pH 7.4, 1 M NaCl, 1 mM EDTA, 0.1% SDS, 1% NP-40), low salt wash buffer (50 mM HEPES pH 7.4, 150 mM NaCl, 1 mM EDTA, 0.1% SDS, 1% NP-40), and PNK buffer (50 mM Tris pH 7.4, 10 mM MgCl_2_, 0.5% NP-40). The RNA-protein complexes were then subjected to 3’ adapter ligation and reverse transcription with SuperScript III Reverse Transcriptase (Invitrogen). The cDNA was purified with Agencourt AMPure XP beads (Beckman Coulter) and subjected to PCR amplification with barcoded primers. The amplified cDNA was purified with Agencourt AMPure XP beads and subjected to high-throughput sequencing on an Illumina NovaSeq platform.

### Skipper windows

eCLIP libraries processed with Skipper (https://github.com/YeoLab/skipper/) (Boyle et al. 2023). Briefly, reads were adapter trimmed with skewer (Jiang et al. 2014) (https://github.com/relipmoc/skewer) and reads less than 20 bp were discarded. UMI were extracted using fastp 0.11.5 (https://github.com/OpenGene/fastp) (Chen et al. 2018). Mapping was then performed against the full human genome (hg38) including a database of splice junctions with STAR (v 2.7.6) (Dobin et al. 2013), allowing up to 100 multimapped regions. When multimapping occurred, reads are randomly assigned to any of the matching position. Reads were then PCR deduplicated using UMIcollaps (https://github.com/Daniel-Liu-c0deb0t/UMICollapse). Enriched “windows” (IP versus SM-Input) was called on deduplicated reads using a GC-bias aware beta-binomial model. Each window (∼100 b.p.) were partitioned from Gencode v38 and associated with a specific type of genomic region (CDS, UTR, proximal introns near splice site… etc). Windows are filtered using FDR < 0.2 and only reproducible windows between two replicates were used.

### Metadensity

To plot enrichment of binding sites across cytoplasmic and whole-cell eCLIPs, we used Metadensity (Her, Boyle, and Yeo 2022) to calculate normalized density. Normalized density is calculated for each nucleotide position in a transcript, density = pi × log_2(pi/qi)_, where pi and qi are the fraction of total reads in IP from either whole-cell or cytoplasmic SNRNP200 libraries respectively that map to nucleotide i. We averaged the relative information over all transcripts with at least one differential enriched window detected by Skipper.

### Immunofluorescence-FISH

Ibidi 18-well μ-slides were pretreated with 0.001% (w/v) poly-D-lysine for 24 h before seeding and reverse transfection. HEK293XT cells were transfected and seeded onto the pretreated 18-well slides at ∼20% confluency. The cells were incubated at 37°C for 48 h. For stressed conditions, sodium arsenite was added to a final concentration of 0.5 mM 45 min before fixation. Cells were fixed by diluting paraformaldehyde to 4% (v/v) directly in the wells and incubating for 1h at 4 °C. The paraformaldehyde was then removed, and the cells were washed three times with dPBS. The cells were then treated with permeabilization solution (0.1% (v/v) Triton X-100 and 1% (w/v) donkey serum in dPBS) for 45 min at 4°C. The samples were washed three times with PBST (0.1% (v/v) Triton X-100 in dPBS). Primary antibodies were added to PBST + 0.1% (w/v) BSA at the following dilutions: α-sc-35 (1:250); α-ERH (1:250); α-G3BP1 (1:1000); α-SNRNP200 (1:100); α-PRPF8 (1:100). The primary antibodies were incubated with the samples for 12 h at 4°C. Following primary antibody treatment, the cells were washed three times with PBST, and the secondary antibodies were added at a 1:2000 dilution in PBST + 0.1% (w/v) PBST for a 2 h incubation at 25°C. The cells were then washed three times with dPBS and fixed again with 4% (v/v) paraformaldehyde for an additional 10 min at 4°C. Following the second fixation, the cells were washed twice with PBST and then treated with FISH-W Buffer (10% (v/v) formamide in 2X SSC) for 5 min at 25°C. After FISH-W Buffer, the poly(A) probe was added to a final concentration of 125 nM in FISH-H Buffer (2 mg/mL BSA + 10% (w/v) dextran sulfate in 2X SSC) and incubated with the cells for 2 h at 37°C. Following probe treatment, the cells were washed twice with FISH-W Buffer and then twice with 2X SSC at 37°C for 5 min. After the final wash, Prolong Diamond with DAPI was added, and the mountant was cured for 12 h at 25°C. The samples were then imaged on an LSM 880 confocal laser scanning microscope in the DAPI, GFP, Cy3, and Cy5 channels with the same laser power and digital gain for all samples.

### Quantification of IF-FISH Images

Fluorescence microscopy images were imported into Fiji for processing and analysis. The number of SC35 foci was quantified by setting the same threshold for the sc-35 channel of each image and using the “Analyze Particles” function to determine the number of speckles per cell. The segmentation function "Watershed" was used to bisect overlapping nuclear speckles for counting. Foci below 0.2 μm^2^ and above 3.0 μm^2^ were filtered from the results. Stress granules were quantified using the same pipeline in the G3BP1 channel but with the size cutoffs of 0.2 μm^2^ and 5.0 μm^2^. The average poly(A) RNA intensity for each image was determined by setting a minimum threshold for poly(A) detection in each image and averaging the intensity of poly(A) RNA that meet this threshold. Intensity profiles for individual images were generated using the “Plot Profile” function in Fiji, and the intensity values were normalized from 0-1 for each fluorescence channel. The average nuclear poly(A) signal intensity was calculated above the background, and then the intensity of the pixels that overlapped with the Dapi signal was averaged (Fiji).

