## Supplementary figures and images for "Large-scale map of RNA binding protein interactomes across the mRNA life-cycle"

### Supplemental Figure 1

Supplemental Figure 1

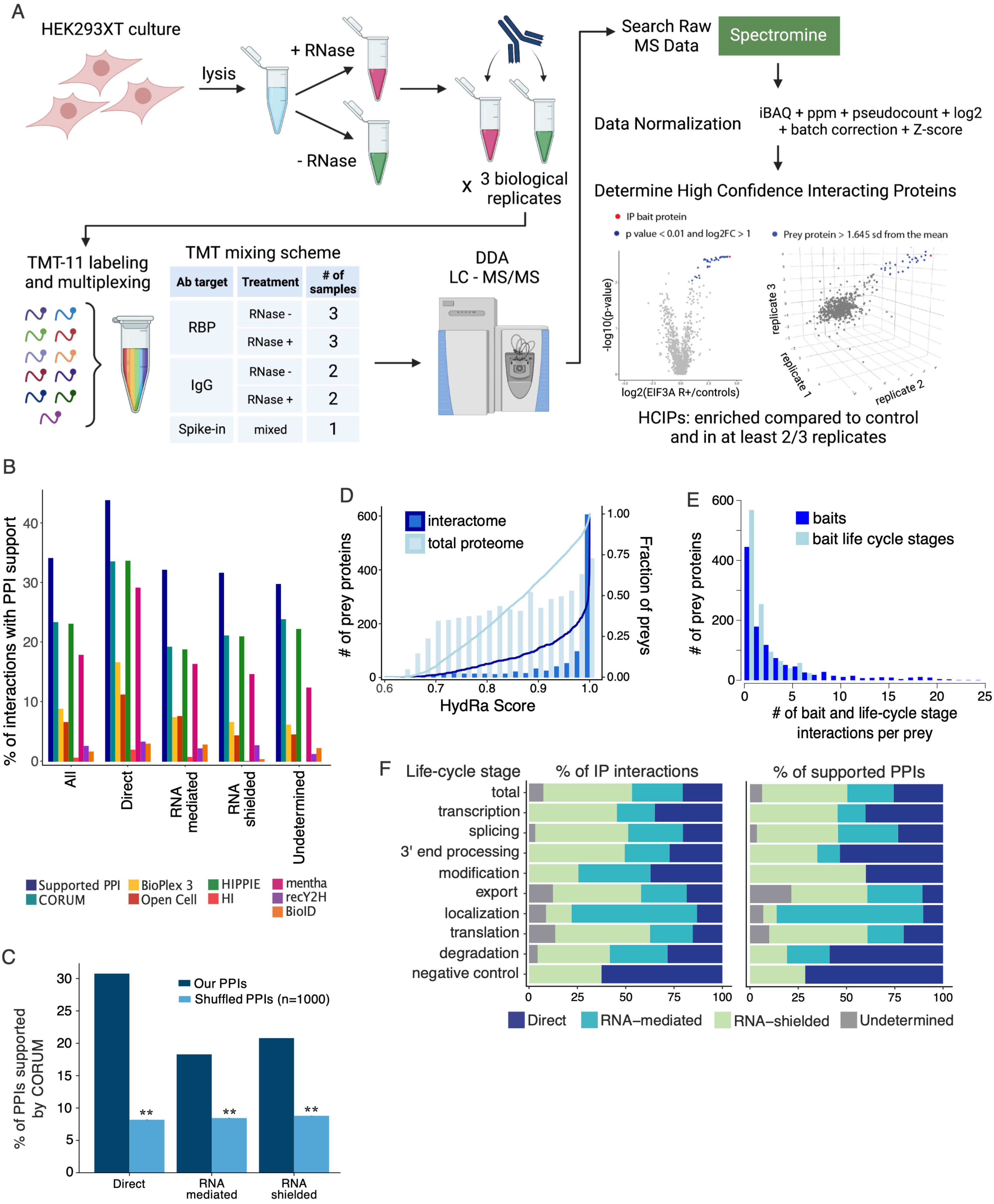

### Supplemental Figure 2

Supplemental Figure 2

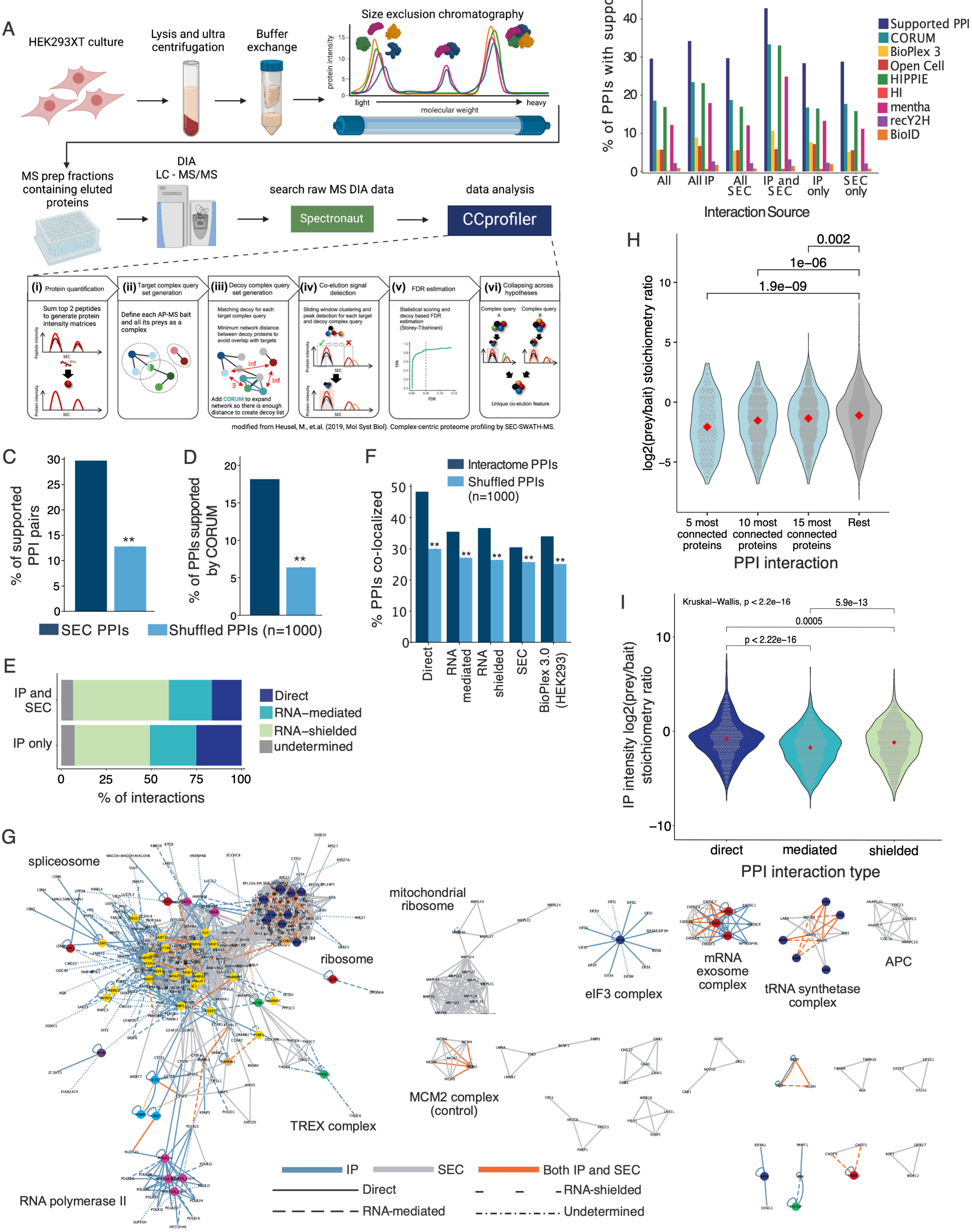

### Supplemental Figure 3

Supplemental Figure 3

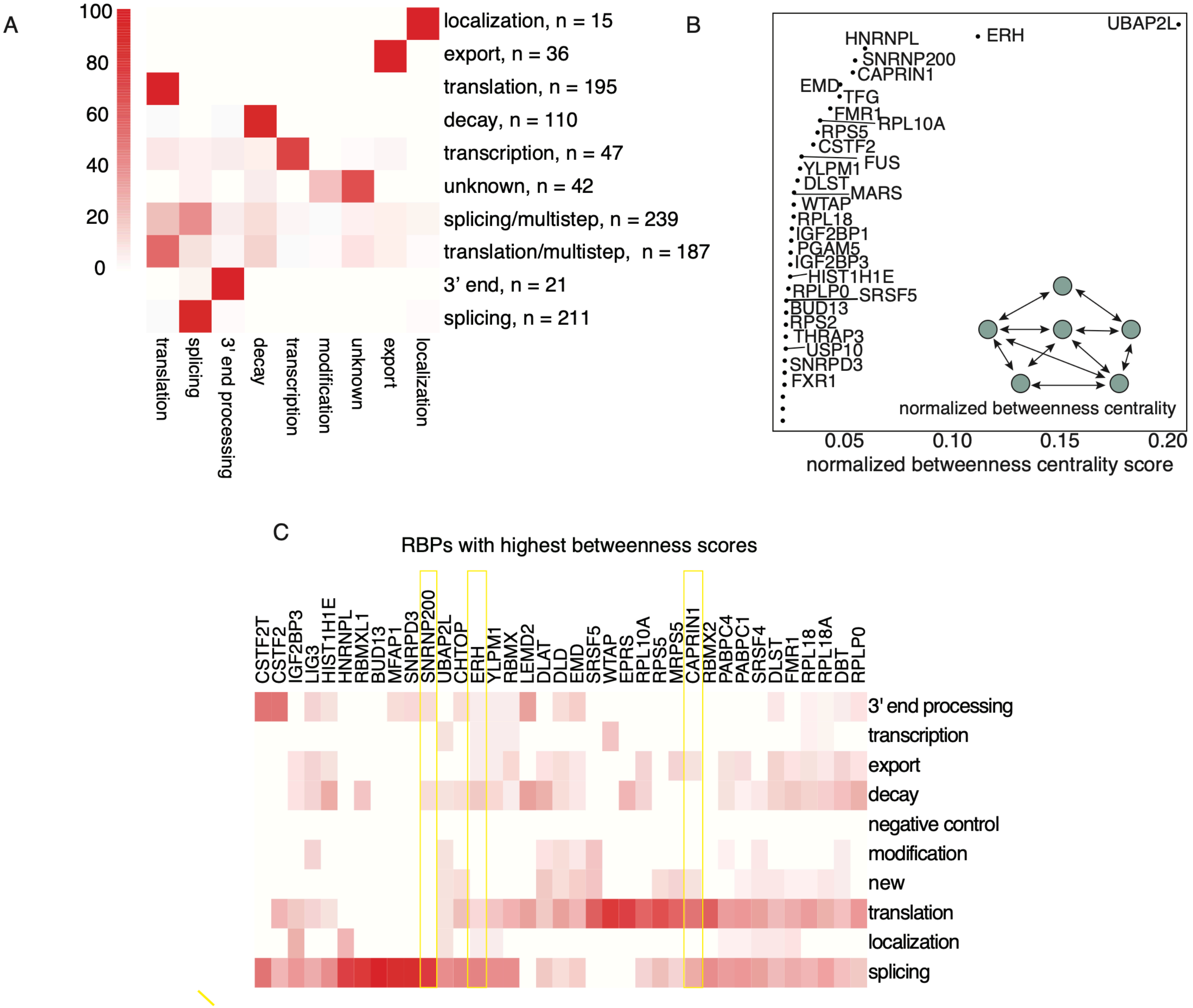

### Supplemental Figure 4

Supplemental Figure 4

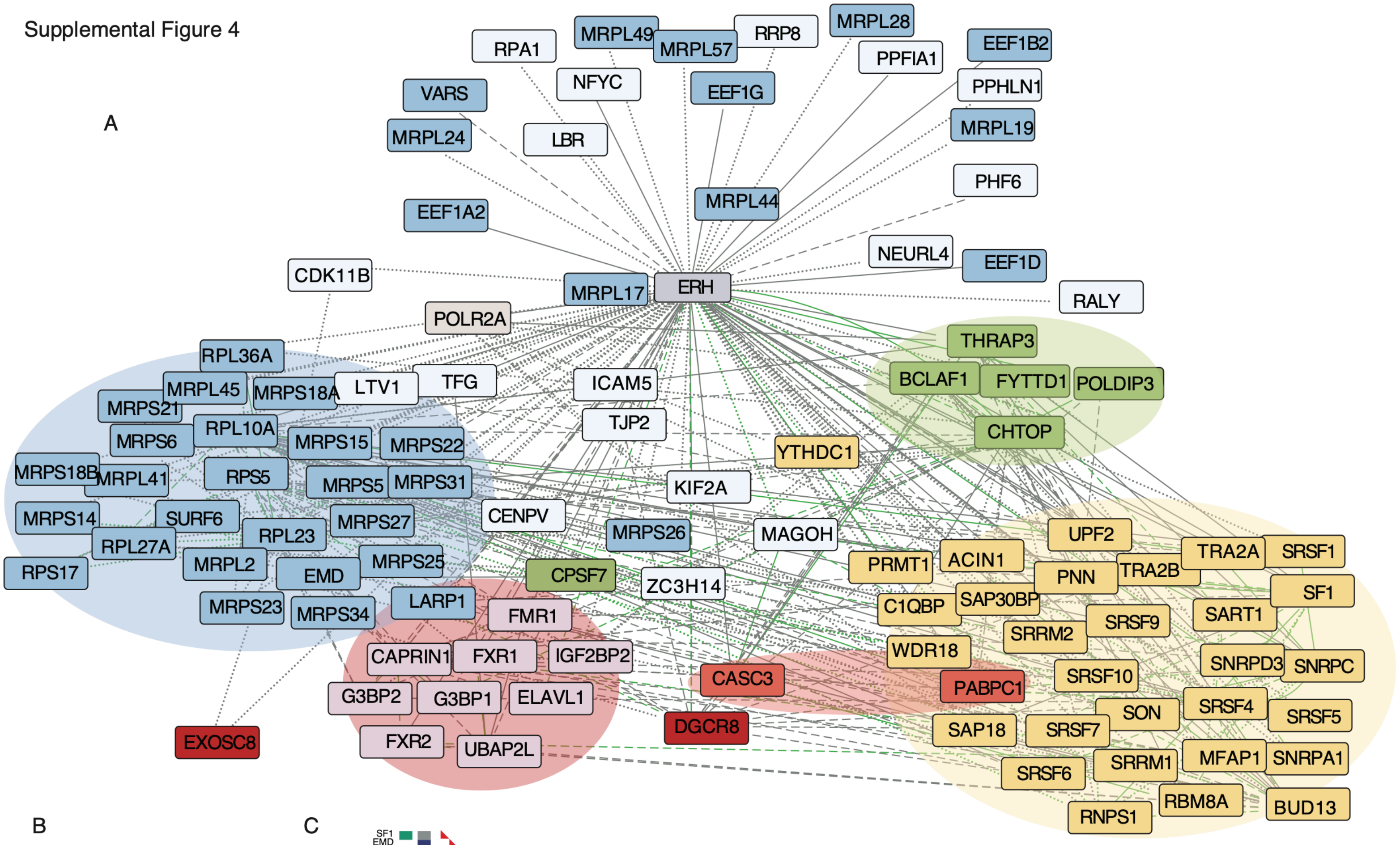


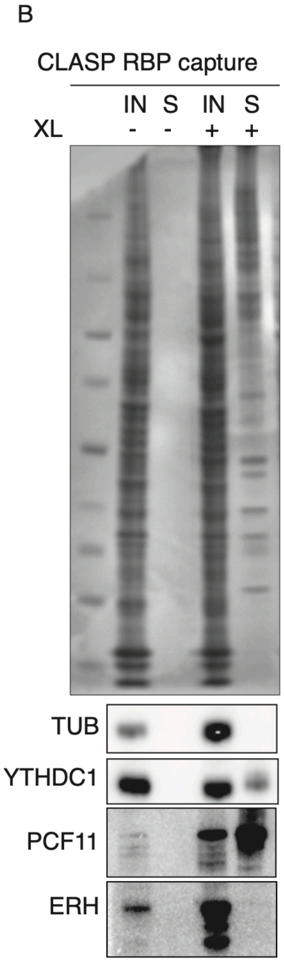


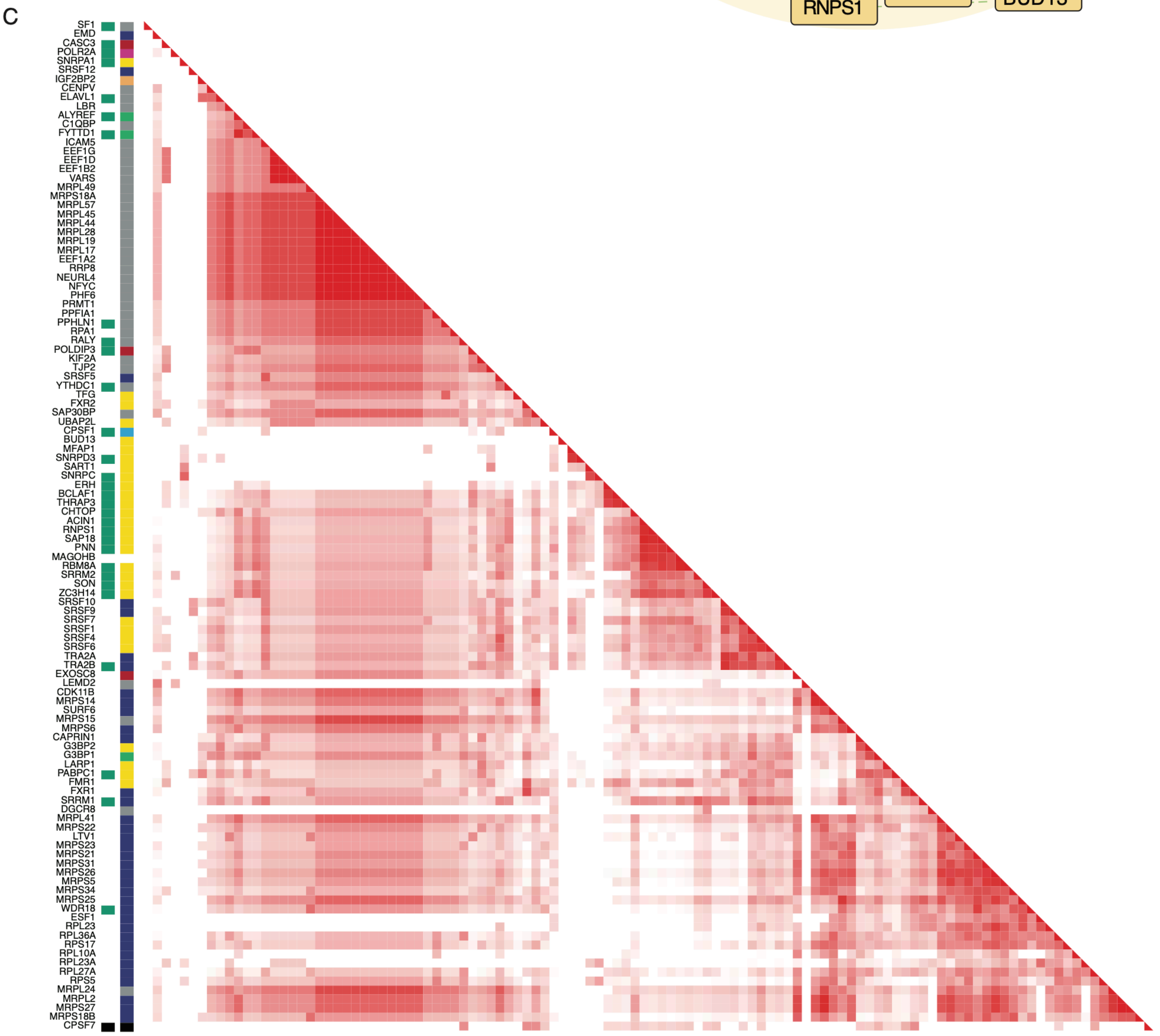

### Supplemental Figure 5

Supplemental Figure 5

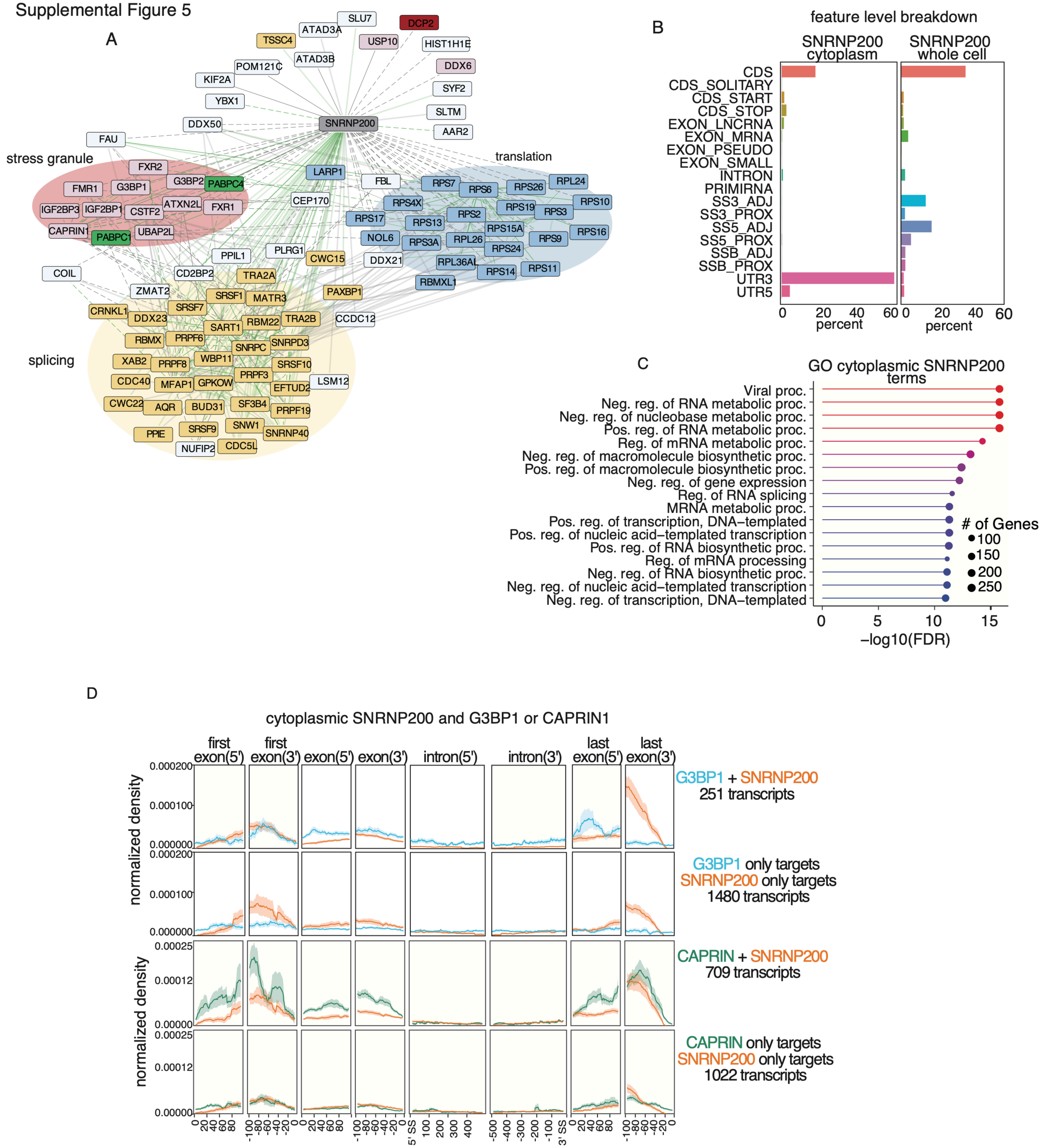

### Supplemental Figure 6

Supplemental Figure 6

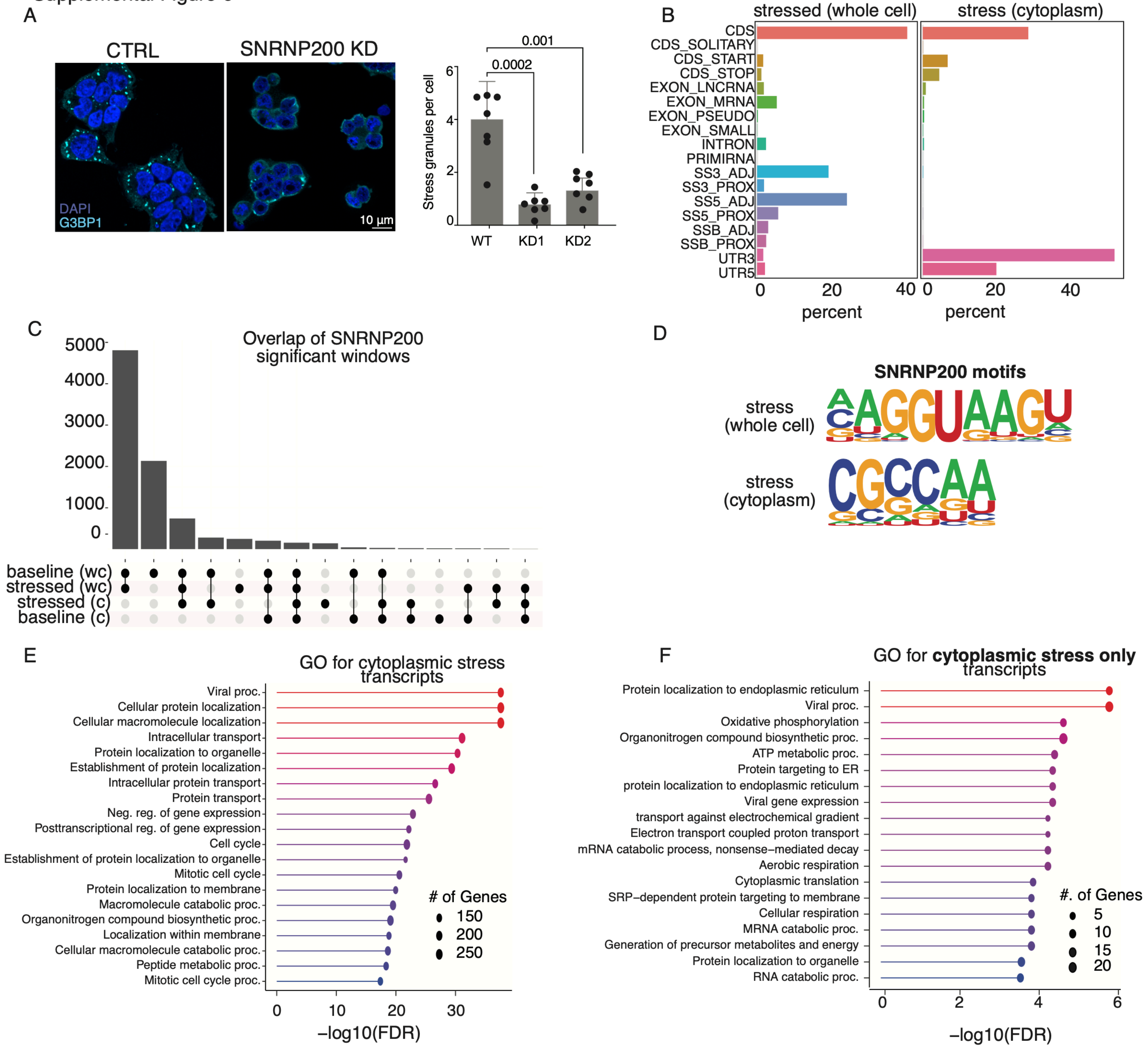
